# A pan-cancer single-cell atlas of pericytes

**DOI:** 10.64898/2026.07.29.741412

**Authors:** Ane Martinez-Larrinaga, Sandra Camargo, Marta Saraiva, Sergi Cervilla, Saioa Garcia-Longarte, Anabel Martinez-Romero, Leonor Gouveia, Hielke van Splunder, Rosa A. Barbella-Aponte, Eva Musulen, Verónica Rodilla, Eva M Galán-Moya, Eduard Porta-Pardo, Arkaitz Carracedo, Isabel Mendizabal, Mariona Graupera

**Author notes:** Joint last authors and authors for correspondence: I.M. and M.G.

## Abstract

Pericytes display marked tissue-specific transcriptional identities, raising the question of whether tumour-associated pericytes converge towards shared adaptive states across cancers. Here, we built a pan-cancer single-cell RNA sequencing atlas of pericytes, integrating nearly four million cells across nine tissues, complemented by spatial transcriptomic analyses. Despite their physiological diversity, pericytes were recurrently expanded in tumours and converged on a common transcriptional program, the tumour-associated pericyte signature (TAPS). TAPS robustly identified pericytes across datasets, outperforming canonical markers in tumour contexts. Tumour-associated pericytes further diversified into specialised states including extracellular matrix (ECM)-associated and interferon (IFN)-responsive programs, which occupy mutually exclusive tumour ecosystems. ECM-associated pericytes were enriched in desmoplastic, fibroblast-rich regions and were associated with adverse clinical outcomes across multiple cancer types. IFN-responsive pericytes accumulated in inflammatory niches, with macrophages implicated as candidate drivers of specialisation. Together, our multi-layered analysis defines convergent and specialised tumour-associated pericyte programs across human cancers.

## Introduction

The tumour microenvironment (TME) is a complex ecosystem composed of malignant cells, stromal populations, immune infiltrates and ECM components that collectively shape disease progression and therapeutic response ^1,2^. Within this dynamic environment, specialised multicellular niches coordinate local cellular interactions and integrate diverse biological signals to regulate tumour behaviour. Recent spatial transcriptomic studies have further shown that multicellular ecosystems are spatially organised within tumours, revealing reproducible vascular and stromal neighbourhoods associated with distinct biological functions^3^. Among these, the perivascular niche (PVN) has emerged as a central organisational hub that orchestrates vascular, stromal and immune processes^4^. Initially defined as the microenvironment surrounding blood vessels and composed primarily of endothelial and mural cells, the concept has progressively expanded to encompass fibroblasts, immune populations and the surrounding ECM. Through the coordinated activity of these cellular and acellular components, the PVN regulates angiogenesis, immune cell trafficking, tissue remodelling and therapeutic responsiveness, thereby exerting a broad influence on tumour evolution^4^. Yet, the cellular composition and organisation of the PVN remain poorly understood across human cancers.

Endothelial cells (ECs) line the inner surface of blood vessels and are central regulators of tumour angiogenesis and immune cell trafficking within the TME^5–8^. Closely associated with ECs are mural cells, comprising vascular smooth muscle cells (vSMCs) and pericytes, which provide structural support to the vascular wall and regulate vessel stabilisation and integrity^9–11^. While vSMCs predominantly surround larger arteries and veins, pericytes closely associate with capillaries. Identification of pericytes has traditionally relied on perivascular localisation together with the expression of markers such as *PDGFRB*, *NOTCH3*, *KCNJ8* and *ABCC9*^9,12,13^. Yet, there is no universal marker that reliably identifies pericytes across tissues. Consistent with this, recent single-cell RNA sequencing (scRNA-seq) atlases have demonstrated pronounced tissue-specific heterogeneity among pericytes and other vascular mural cells across human organs, highlighting the strong influence of tissue context on pericyte identity^11,14,15^. In tumours, pericytes undergo profound phenotypic remodelling, characterised by abnormal vessel coverage, impaired vascular interactions, and the acquisition of transcriptional programs associated with inflammation, ECM remodelling, and loss of pericyte identity^16–19^. These observations raise the fundamental question of whether tumour-associated pericytes preserve their physiological organotypic specialisation or instead converge towards shared tumour-associated states across tissues.

scRNA-seq has transformed our understanding of the TME by enabling the unbiased characterisation of cellular heterogeneity and the identification of specialised cellular states^20–22^. These approaches have generated comprehensive atlases of immune, stromal, and vascular populations, revealing diverse states of T-cell dysfunction, macrophage polarisation, fibroblast specialisation, and EC activation across human cancers^23–26^. Although recent vascular atlases have substantially advanced the molecular definition of EC and mural populations, mural cells remain comparatively underrepresented in tumour-focused single-cell studies. Their low abundance, inefficient recovery during tissue dissociation, and the absence of definitive markers have hindered their systematic study, often resulting in low-resolution annotation or their inclusion within broader vascular compartments^11,27,28^. As a result, a comprehensive understanding of pericyte and vSMC identities, heterogeneity, and remodelling in human cancers remains poorly understood.

Here, we built a pan-cancer single-cell atlas of the PVN across multiple human tumour types to systematically characterise pericyte identity, define tumour-associated transcriptional programs, and investigate specialised pericyte states. By integrating scRNA-seq and spatial profiling approaches, we establish a transcriptional reference for pericyte identification, uncover convergent tumour-associated pericyte states, and reveal spatially organised pericyte ecosystems associated with ECM remodelling and immune regulation. Together, our findings provide a comprehensive resource for understanding pericyte biology in cancer and offer new insights into how the PVN shapes the organisation of the tumour ecosystem.

## Results

### A pan-cancer atlas reveals selective expansion of pericytes

To generate a comprehensive pan-cancer reference of the PVN, we compiled 56 publicly available scRNA-seq atlas studies spanning nine human tissues (**Fig. 1a,b**). In total, 296 non-tumour and 615 tumour patient specimens were included, prioritising primary tumours, tumour-adjacent tissues, and matched non-tumour samples, while excluding treated or metastatic cases (**Fig. 1c**). To ensure technical consistency, only datasets generated using droplet-based platforms (e.g., 10x Genomics) were retained (**Table S1**). For each tissue, datasets were processed independently, including quality control, donor-level batch correction using Harmony, and graph-based clustering (**Extended Data Fig. 1a**). Major cellular compartments were identified and annotated based on canonical marker expression (**Table S2**). These comprised epithelial (*EPCAM*, *KRT8/18*), immune cells, further stratified into lymphoid (*CD4*, *CD8A*, *CD3G*, *CD79A/B*, *NKG7*) and myeloid (*C1QA*, *C1QB*, *CD68*) lineages, ECs (*PECAM1*, *CDH5*), fibroblast (*PDGFRA*, *COL1A1*), mural cells (*PDGFRB*, *RGS5*, *ACTA2*), and dividing (*MKI67*, *TOP2A*) populations (**Table S3**). In total, nearly four million cells were analysed, providing a comprehensive representation of the TME across nine tissues (**Extended Data Fig. 1b**). These cell types were further grouped into broader compartments, including epithelial, immune, proliferative, and the PVN (**Fig. 1d**). Overall, 527,857 cells (13.6%) were classified as part of the PVN, comprising ECs, fibroblasts, and mural cells (**Fig. 1e**).

**Figure 1.**
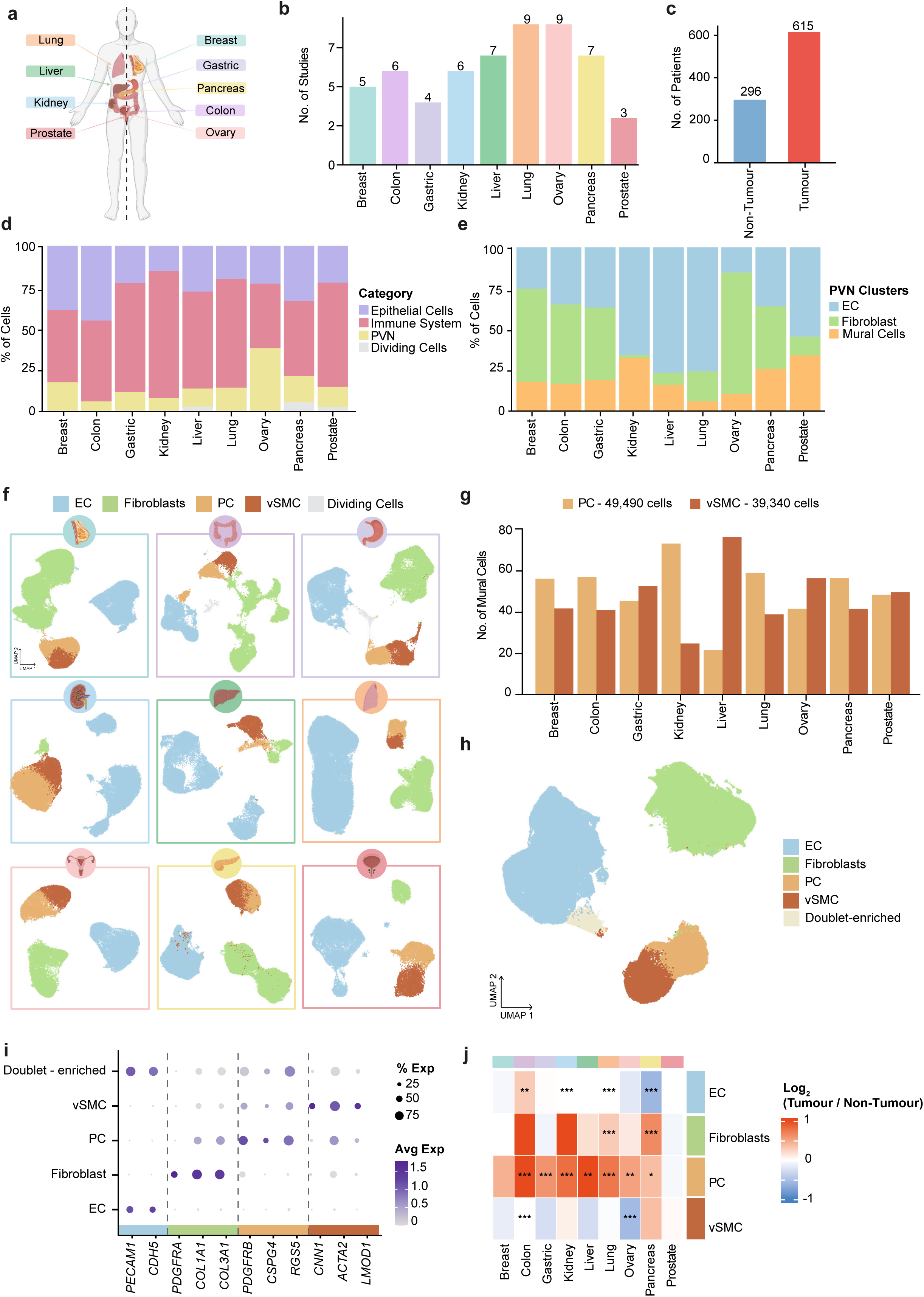
A pan-cancer atlas reveals selective expansion of pericytes. **a)** Diagram illustrating the tissues analyzed: breast, colon, gastric, kidney, lung, liver, ovary, pancreas, and prostate. **b)** Bar plot representing the number of publications included for each tissue. **c)** Bar plot representing the number of patients per condition; non-tumour (blue), and tumour samples (red). **d)** Stacked bar plot showing the cellular composition (percentage) of epithelial (purple), immune (pink), perivascular (yellow), and proliferating (grey) compartments across all tissues. **e)** Stacked bar plot showing the percentage of PVN populations: ECs (blue), fibroblasts (green), and mural cells (orange) across all tissues. **f**) UMAP plots showing the identification of the main PVN subtypes per tissue, organised alphabetically: breast, colon, gastric, kidney, liver, lung, ovary, pancreas, and prostate. Cell populations are annotated as EC, fibroblast, pericytes (PC) and vSMC. **g)** Bar plot showing the number of PC and vSMC across all tissues. **h)** UMAP plot of the integrated Seurat object containing all PVN cells across all tissues. **i**) Dot plot showing canonical markers for the main PVN populations. The size of the dots represents the percentage of cells expressing the marker, and the colour intensity reflects the average expression level across the identified PVN cell types. **j)** Heatmap showing the log_2_FC in cell proportions between tumour and non-tumour samples estimated by propeller. Positive and negative values indicate expanded and contracted populations in tumours, respectively. Statistical significance was assessed using an empirical Bayes moderated t-test, with adjusted P values indicated as: P ≤ 0.05 (*), P ≤ 0.01 (), and P ≤ 0.001 (*). Tissue colour scheme as in **a**.

To enable high-resolution analysis, PVN cell types were subsetted and re-analysed independently within each tissue (**Fig. 1f**). This approach enabled clear discrimination of the two canonical mural cell populations: pericytes (*PDGFRB*, *RGS5*, *NOTCH3*) and vSMCs (*ACTA2*, *CNN1*, *LMOD1*) (**Extended Data Fig. 1c**). We identified a total of 49,490 pericytes and 39,340 vSMCs (**Fig. 1g**). The complete perivascular compartment from each tissue was subsequently integrated, and re-annotated using canonical markers (**Fig. 1h,i** and **Table S2**). Within this integrated object, ECs accounted for 42.9%, fibroblasts for 35.9%, pericytes for 11.6%, and vSMCs for 9.2% (**Extended Data Fig. 1d**). A minor population enriched for predicted doublets was excluded from downstream analyses **(Extended Data Fig. 1e-h)**. This cluster lacked a coherent marker gene profile consistent with any single PVN cell type (**Extended Data Fig. 1i**). Differential abundance analysis using propeller^29^, revealed that pericytes were significantly enriched in tumours across seven of nine tissues, representing the most consistent expansion among perivascular populations (**Fig. 1j** and **Extended Data Fig. 1j**).

### A convergent tumour-associated transcriptional program defines pericyte identity

Next, we sought to deconstruct the transcriptional programs underlying PVN cell type identity. Tumour and non-tumour samples were analysed separately by comparing each cell type with the remaining perivascular cells within the same tissue. Results for ovarian and breast are shown as representative examples in **Fig. 2a** and **Extended Data Fig. 2a**, respectively, whereas results from other tissues are provided in **Table S4**. By intersecting genes consistently upregulated across all nine tissues, we defined core transcriptional signatures for ECs, fibroblasts, pericytes and vSMCs (**Extended Data Fig. 2b** and **Table S5**). A larger subset of genes was shared across tissues in tumour than in non-tumour samples in all PVN populations (**Fig. 2b**). The difference was most pronounced in pericytes: only four genes were shared in non-tumour samples, expanding to 53 genes in tumours (**Fig 2b,c** and **Extended Data Fig. 2b**). We therefore define this set of 53 genes as the tumour-associated pericyte signature (TAPS), a convergent pan-cancer transcriptional program (**Fig. 2d**). Notably, 46 out of 53 TAPS genes showed higher expression in tumour than non-tumour pericytes (**Extended Data Fig. 2c**), while some TAPS genes were also expressed in non-tumour samples (**Extended Data Fig. 2d**). To assess whether the increased transcriptional overlap in tumours could be explained by unequal cell numbers between tumour and non-tumour groups, we performed a downsampling analysis in which tumour cells were randomly subsampled to match the number of non-tumour cells within each tissue (**Extended Data Fig. 2e**). The number of shared genes remained consistently higher in tumours across all PVN populations (**Extended Data Fig. 2f**, empirical P < 0.001). Collectively, these findings indicate the emergence of a convergent tumour-associated pericyte state regardless of tissue of origin.

**Figure 2.**
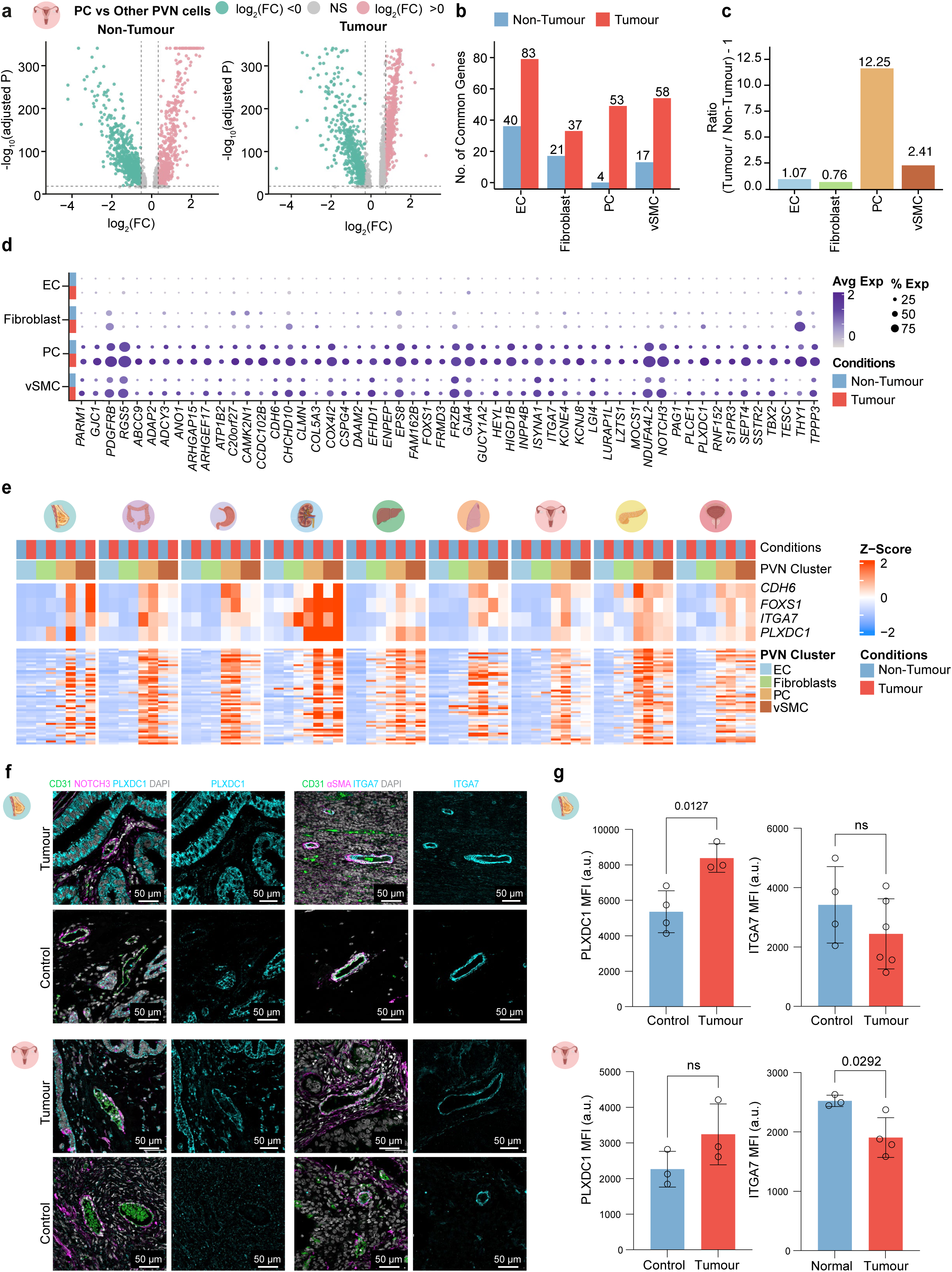
A convergent tumour-associated transcriptional program defines pericyte identity. **a)** Volcano plots of DEGs in pericytes (PC) versus other PVN populations, in the ovary dataset. Comparison between non-tumour (left) and tumour (right) samples. Pink and green dots represent significantly upregulated and downregulated genes, respectively; grey dots (NS) indicate non-significantly expressed genes. **b)** Bar plot representing the number of genes identified in both non-tumour and tumour samples across all studied tissues, per PVN subtypes. **c)** Bar plot showing the ratio of shared genes between tumour and non-tumour conditions for each PVN cell type, expressed as (Tumour/Non-Tumour) - 1, where 0 indicates equal sharing across conditions. Positive values indicate a greater number of shared genes in tumour than non-tumour samples. **d)** Dot plot showing the average expression (colour scale) and percentage of expression (dot size) of the 53 genes defining the tumour-associated pericyte signature (TAPS) across the different PVN compartments. **e)** Heatmap showing the average TAPS expression (Z-score) across cell subsets in non-tumour (blue) and tumour (red) conditions. The upper panel highlights the four genes selected for immunofluorescence validation (CDH6, FOXS1, ITGA7, PLXDC1) at expanded scale; the lower panel shows the remaining TAPS genes. Tissues are ordered alphabetically (breast, colon, gastric, kidney, liver, lung, ovary, pancreas, and prostate). **f)** Representative immunofluorescence images of tumour and control human breast (top) and ovarian (bottom) tissues showing CD31+ blood vessels (green) surrounded by NOTCH3+ or αSMA+ mural cells (magenta), together with single TAPS-markers PLXDC1 and ITGA7 (cyan). **g)** Quantification of PLXDC1 and ITGA7 mean fluorescence intensity (MFI) in breast and ovarian tissues. Each dot represents one patient. Data are shown as mean ± s.e.m. Statistical significance was assessed using the Mann-Whitney test; P > 0.05 was considered non-significant (ns).

Next, we investigated whether individual TAPS genes labelled pericytes specifically *in situ*. To prioritise candidate markers, we first examined the expression of all 53 TAPS genes across tissues, stratified by condition and PVN cell type (**Fig. 2e**). We then selected genes that were consistently upregulated in tumours relative to non-tumour tissues, particularly within mural cells, and prioritised candidates previously implicated in tumour-associated stromal niches for which validated commercial antibodies were available. Based on these criteria, we selected plexin domain-containing protein 1 (PLXDC1), integrin α7 (ITGA7), cadherin 6 (CDH6), and the transcriptional factor FOXS1^30–33^. Ovarian and breast tissues were used as representative samples for immunofluorescence validation (**Fig. 2f** and **Extended Data Fig. 2g**). Vascular and perivascular populations were identified by co-staining for CD31 (EC) and αSMA or NOTCH3 (mural cells). The mean fluorescence intensity (MFI) of each candidate marker was subsequently quantified within αSMA or NOTCH3 positive areas (**Fig. 2g** and **Extended Data Fig. 2h**). All four proteins were detected in mural cell populations, however, PLXDC1, FOXS1 and CDH6 were also detected in other cell populations in both tumour and control tissues (**Fig. 2f** and **Extended Data Fig. 2g**). In contrast, ITGA7 expression was largely restricted to mural cells, where it mainly overlapped with αSMA-positive vessel-associated cells (**Fig. 2f**). PLXDC1 showed significantly higher expression in breast tumours, and a trend towards increased expression in ovarian tumours. A similar pattern was observed for CDH6 in breast tumours (**Fig. 2g** and **Extended Data Fig. 2h**). While these analyses identify PLXDC1 as a tumour-enriched mural cell marker, no single candidate reliably distinguished pericytes from vSMCs or discriminated tumour-associated from non-tumour pericytes.

### TAPS enables robust pericyte identification across datasets and spatial contexts

The limited discriminatory power of individual TAPS genes prompted us to investigate whether their combined expression could serve as a robust transcriptional signature to identify pericytes. TAPS activity was consistently higher in pericytes relative to vSMCs, fibroblasts, and ECs in both the integrated and individual tissue datasets (**Fig. 3a** and **Table S6**). To validate these findings in an independent context, we analysed a scRNA-seq dataset of human brain tumours from *Wälchli et al*. dataset^34^, a cancer type not included in our discovery cohort. Consistent with our findings, TAPS activity was also elevated in pericytes relative to other PVN cell types in this independent dataset (**Fig. 3b** and **Extended Data Fig. 3a,b**).

**Figure 3.**
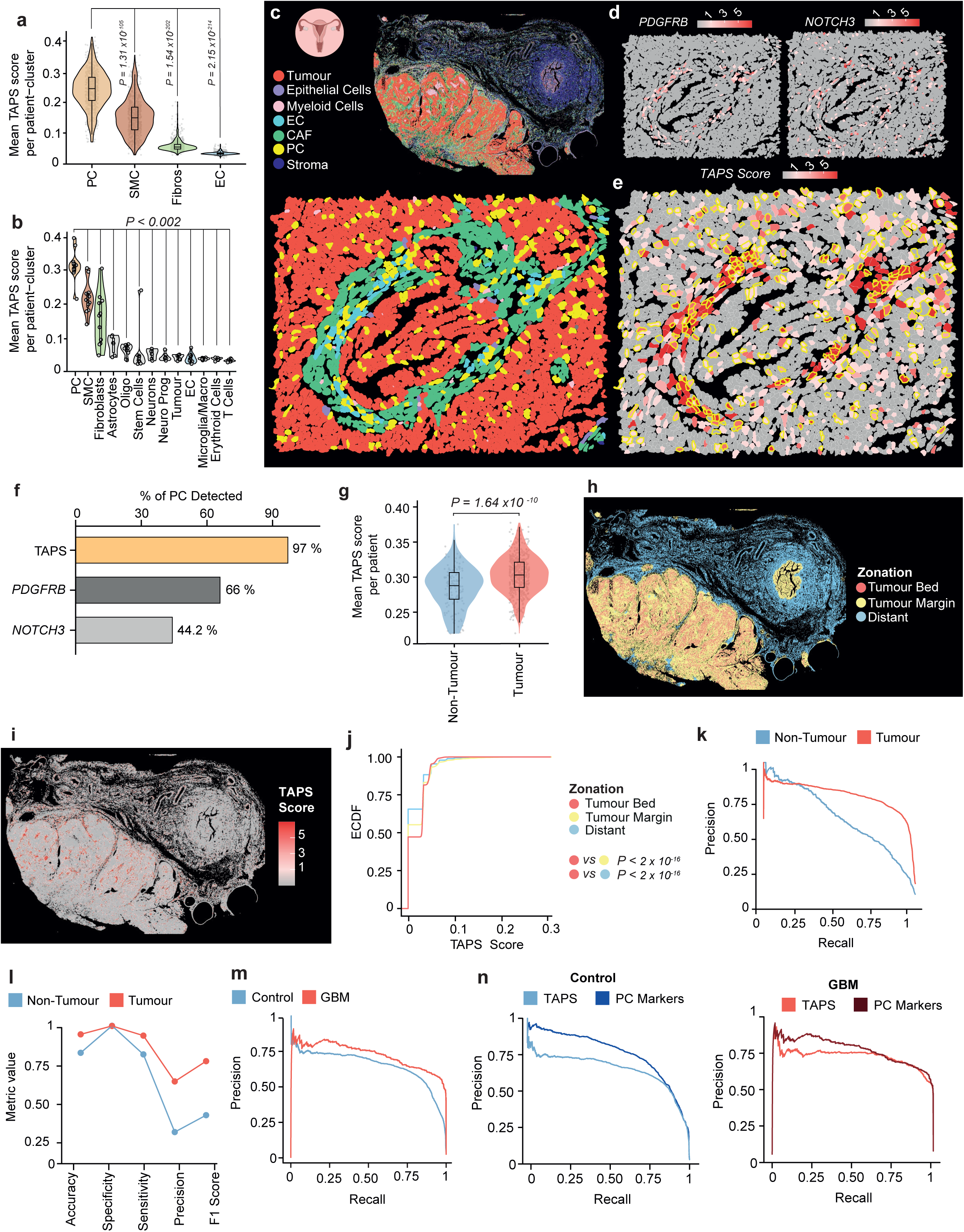
TAPS enables robust pericyte identification across datasets and spatial contexts. **a)** Violin plots showing per-patient mean TAPS score for each PVN cell-type. Statistics were calculated using pairwise Wilcoxon rank-sum test comparing pericytes (PC) against all other PVN cell types. **b)** Violin plots showing per-patient mean TAPS score for each PVN cell-type based on the clusters defined in *Wälchli et al.* Statistics were calculated using pairwise Wilcoxon rank-sum test comparing PC against all other clusters. The displayed P-value represents the largest observed value. Abbreviations: Oligo, oligodendrocytes; Neuro prog, neuronal progenitors; Mac, macrophages. **c)** Spatial distribution of annotated cell types in ovary. Representative images show the full tissue section (top) and a magnified inset (bottom). **d)** Representative images of the spatial expression of *PDGFRB* and *NOTCH3*. **e)** Representative image of the spatial distribution of the TAPS score; the yellow border represents PC identified in panel **c**. **f)** Bar plot showing the percentage of PC identified by TAPS compared to the annotations provided in the original ovary dataset. **g)** Violin plots showing the mean TAPS score per patient within the PC cluster, comparing non-tumour and tumour samples. Statistics were calculated using a Wilcoxon rank-sum test. **h)** Spatial distribution of the different tissue zones: tumour bed (red), tumour margin (yellow), and distant tissue (blue), in the breast sample**. i)** Spatial distribution of the TAPS UCells score across the ovary sample. **j)** Empirical cumulative distribution function (ECDF) curves showing TAPS UCell scores across tumour zonation regions in ovary. Statistical significance was estimated using the Kolmogorov-Smirnov test. **k)** Precision-recall curves of the model used in the discovery dataset to distinguish pericytes in non-tumour and tumour samples. **l)** Dot plot showing metrics from the confusion matrix of the discovery dataset to distinguish PC in non-tumour and tumour samples. **m)** Precision-recall curves of the model applied to the independent dataset to distinguish PC in control and tumour samples. Abbreviations: GMB, glioblastoma. **n)** Precision-recall curves comparing the discriminatory performance of TAPS and canonical pericyte marker signature scores for pericyte identification in the independent validation cohort under control (left) and tumour conditions (GBM, right).

We next evaluated the performance of TAPS in spatial transcriptomic data using publicly available Xenium (10x Genomics) datasets from ovarian and breast tumours (**Table S1**). Throughout the manuscript, analyses from the ovarian tumour dataset are shown in the main figures, whereas corresponding analyses from an independent breast tumour dataset are presented in the **Extended Data Figures**. Following preprocessing and initial cell-type annotation using the same marker-based annotation strategy as in the scRNA-seq atlas, pericytes were identified for downstream analyses (**Fig. 3c**, **Extended Data Fig. 3c,** and **Table S2**). Evaluation of single canonical pericyte markers *PDGFRB* and *NOTCH3* revealed sparse and fragmented expression patterns, consistent with the limited transcript detection and transcript drop-out characteristic of targeted spatial transcriptomic technologies such as Xenium, which often hinder accurate discrimination between pericytes and vSMCs (**Fig. 3d** and **Extended Data Fig. 3d**). In contrast, projection of TAPS activity enabled robust identification of pericyte populations across both tumour types (**Fig. 3e** and **Extended Data Fig. 3e**). Using the published cell type annotations from the ovarian spatial transcriptomic dataset as the reference, high TAPS activity was detected in 97% of pericytes, whereas high *PDGFRB* and *NOTCH3* expression was observed in only 66% and 44.2% of these cells, respectively (**Fig. 3f** and **Extended Data Fig. 3f**). These findings demonstrate that TAPS provides a more sensitive and robust approach for pericyte identification in spatial transcriptomic datasets.

Having established that TAPS robustly identifies pericytes across datasets, we next investigated whether TAPS activity reflects the TME. Indeed, TAPS activity was consistently elevated in tumour-derived pericytes relative to non-tumour counterparts (**Fig. 3g** and **Table S7**). In the *Wälchli et al*. dataset^34^, TAPS activity showed a similar pattern, with higher scores in tumour-associated pericytes, reaching statistical significance at the single-cell level (P = 1.08 x10^-80^) but not in patient-level pseudobulk analysis (P = 0.0855), likely reflecting the limited donor number in this dataset (**Extended Data Fig. 3g,h**). To determine whether this tumour-associated elevation of TAPS activity could also be observed *in situ*, samples were segmented using distance-based analysis into tumour bed (colocalised with cancer cells), tumour margin (<100 μm from cancer cells), and distant regions (>100 μm) (**Fig. 3h** and **Extended Data Fig. 3i**). Consistent with the single-cell analyses, TAPS activity increased progressively from distant to tumour-margin to tumour-bed regions (**Fig. 3i,j** and **Extended Data Fig. 3j,k**). Together, these findings indicate that TAPS captures convergent tumour-associated pericyte transcriptional programs across tissues and experimental platforms.

While TAPS activity scores consistently distinguished pericytes across datasets, we next formally quantified this discriminative power using logistic regression models trained independently on tumour and non-tumour samples. The TAPS-based classifier accurately distinguished pericytes from non-pericyte PVN populations (*i.e.,* ECs, fibroblasts and vSMCs) in both conditions, with higher performance in tumour samples (**Fig. 3k**; AUPRC non-tumour = 0.605, tumour = 0.764). Precision and F1 scores were also higher in tumour-derived samples while maintaining high sensitivity in both conditions (**Fig. 3l**), indicating that TAPS performance improves in the TME. Application of the pre-trained model to the independent *Wälchli et al*. dataset^34^ similarly yielded robust pericyte identification (**Fig. 3m**; AUPRC non-tumour = 0.652, tumour = 0.725). Benchmarking against canonical marker-based signature (*ABCC9*, *ANPEP*, *CD248*, *CSPG4*, *HIGD1B*, *KCNJ8*, *NDUFA4L2*, *PDGFRB*, *RGS5*, and *NOTCH3*)^13^ on the same dataset (**Extended Data Fig. 3l**) showed superior canonical marker performance in non-tumour tissue (**Fig. 3n**; AUPRC canonical = 0.761, TAPS = 0.652), whereas TAPS outperformed canonical markers in tumour (**Fig. 3n**; AUPRC TAPS = 0.725, canonical = 0.677), indicating that TAPS more accurately discriminates tumour-associated pericytes than canonical marker combinations.

### Tumour-associated pericytes exhibit distinct functional states

We next investigated the extent of functional heterogeneity within the pericyte compartment. To this end, we subsetted the mural cells from the integrated PVN dataset and re-analysed them using the same integration and clustering workflow applied during the atlas construction. vSMCs were retained as a stable mural reference to distinguish pericyte-specific programs from broader contractile states. Graph-based clustering revealed substantial heterogeneity within the mural compartment, identifying nine pericyte and three vSMC subpopulations (**Extended Data Fig. 4a,b**). Given the limited availability of established markers for mural subtypes, we combined cluster-specific gene signatures with pathway enrichment analyses to define their identities (**Fig. 4a** and **Table S8**). The top differentially expressed genes associated with each cluster are provided in **Table S9**. Among these populations, we identified a quiescent pericyte state enriched for homeostatic and low-metabolic-activity programs and characterised by *HIGD1B* expression, which we defined as basal pericytes (**Fig. 4b**). In contrast, the remaining populations reflected specialised pathological states associated with tumour remodelling. These included ECM-associated pericytes enriched for matrix-remodelling programs and expressing *POSTN* and *PLXDC1*, angiogenic pericytes enriched for vascular developmental pathways and expressing *GUCY1A1*/*GUCY1B1*, and energetic pericytes with elevated oxidative metabolic activity and marked expression of *COX4I2/COX7C*. We also identified two immune-related states characterised by inflammatory (*CCL19*) and interferon-responsive (*ISG15*, *CXCL10*) transcriptional programs linked to leukocyte chemotaxis, myeloid differentiation, IFN-γ signalling, and immune activation (**Fig. 4b**). Notably, several of the identified pericyte states recapitulated transcriptional programs previously associated with tumour-associated mural populations in independent studies^27,28^ (**Extended Data Fig. 4c,d**).

**Figure 4.**
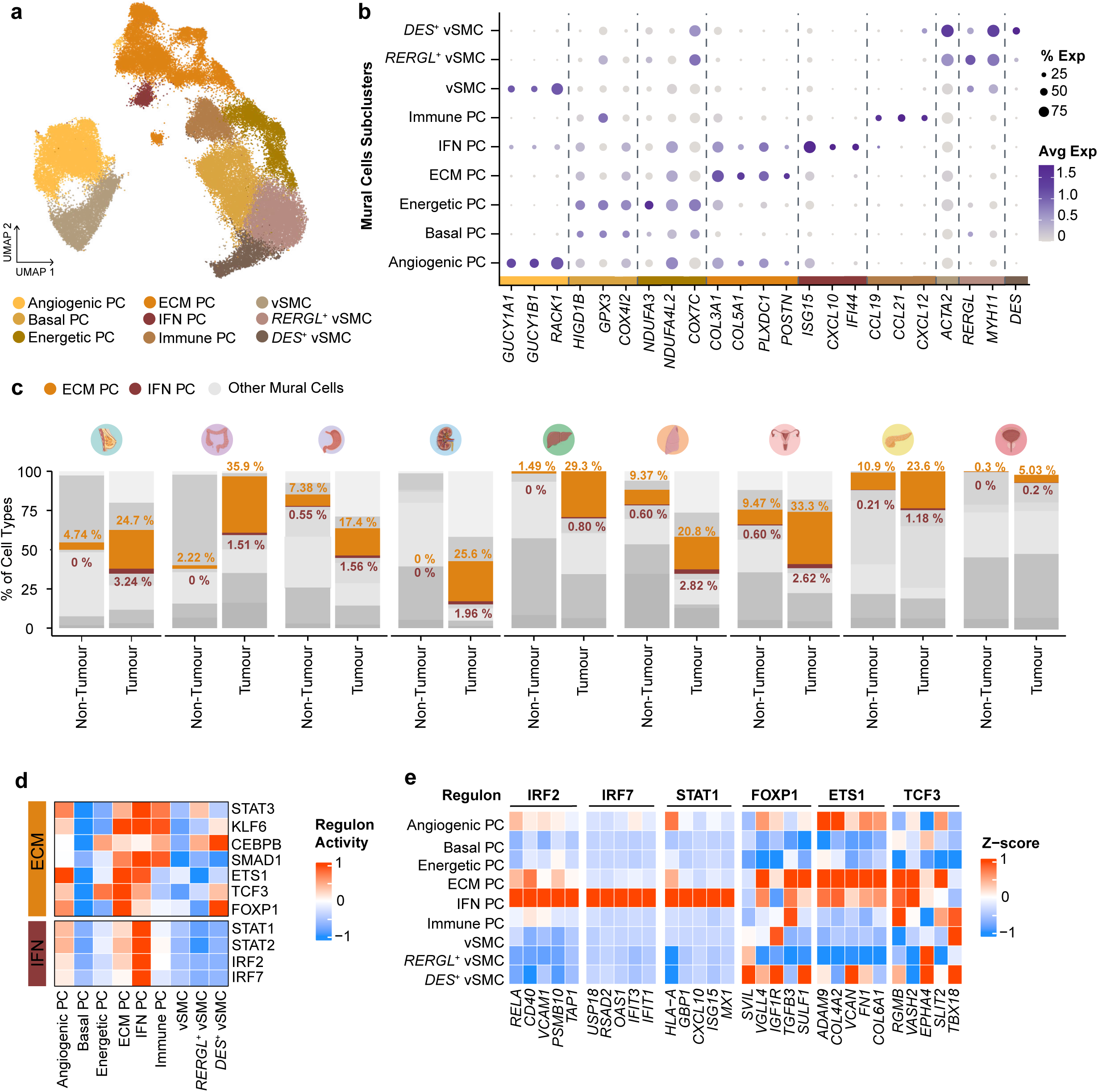
Tumour-associated pericytes exhibit distinct functional states. **a)** UMAP representation of mural cells from the integrated PVN dataset. Cells are coloured according to transcriptionally defined subtypes, including multiple pericyte (PC) states and vSMC populations. **b)** Dot plot showing the average expression (colour intensity) and percentage of expression (dot size) of marker genes across mural cell subtypes. Genes are displayed in rows and cell subtypes in columns. **c)** Stacked bar plots showing distributional changes of mural cell subtypes across non-tumour (left) and tumour (right) samples in nine tissues, organized alphabetically. Colours represent specific PC subtypes: ECM-associated PCs and IFN-responsive PCs. **d)** Heatmap showing regulon activity for the transcription factors (TFs) predicted by SCENIC. Each column represents a mural cell subtype and each row a TF; red indicates higher regulon activity and blue indicates lower activity. All displayed regulons are motif-validated **e)** Heatmap showing target gene expression across mural cell subtypes. Each row represents a transcriptionally defined PC or vSMC subtype, and each column represents a target gene, grouped by the corresponding TF. Expression values are scaled (Z-score) across cell types, with red indicating higher expression and blue lower expression.

We next examined the distribution of these specialised pericyte states across tumour and non-tumour samples using differential abundance analyses using propeller^29^. Basal pericytes were depleted in several tumour types, indicating a loss of homeostatic pericyte programs within the TME (**Extended Data Fig. 4e**). In contrast, ECM-associated and IFN-responsive pericytes were largely absent from non-tumour samples but markedly enriched in tumours. ECM-associated pericytes exhibited broad expansion across tissues, whereas IFN-responsive pericytes displayed a more restricted distribution, being preferentially enriched in breast, kidney, liver, lung, and ovarian tumours (**Fig. 4c**). Notably, ECM-associated pericytes constituted a substantially larger population than IFN-responsive pericytes across all tissues (**Fig. 4c**). The remaining pericyte subtypes did not exhibit consistent patterns of enrichment or depletion (**Extended Data Fig. 4e**).

To further characterise the regulatory networks underlying these specialised pericyte states, we performed SCENIC^35^ analysis, which infers transcription factor regulons and quantifies their coordinated activity at the single-cell level (**Fig. 4d**). IFN-responsive pericytes displayed enrichment of STAT and IRF transcription factor families, such as IRF7 and STAT3, with target genes including canonical immune-effector genes including *CXCL10*, *OAS1*, *MX1*, *IFIT1*, *CD40*, and *VCAM1* (**Fig. 4e**). In contrast, ECM-associated pericytes exhibited a broader regulatory landscape characterised by activation of ETS1-and SMAD1-driven regulons, whose target programs encompassed extracellular matrix-remodelling genes such as *TGFB3*, *FN1*, *COL4A2*, and *ADAM9* (**Fig. 4d,e**). Together, these analyses indicate that tumour-associated pericytes adopt distinct regulatory programs, including programs associated with inflammatory activation and ECM remodelling.

### TAPS enables spatial resolution of specialised tumour-associated pericyte states

We next examined whether specialised pericyte states identified by single-cell transcriptomics could be resolved *in situ*. Given their consistent enrichment across tissues, we focused on the spatial characterisation of ECM-associated and IFN-responsive pericytes. Projection of scRNA-seq derived ECM- (**Fig. 5a**, left) and IFN-associated (**Fig. 5a**, right) transcriptional programs onto spatial transcriptomic datasets from ovarian and breast tumours (**Extended Data Fig. 5a**) revealed broad stromal activity that was not restricted to pericytes, reflecting the shared matrix-remodelling and inflammatory signatures in several stromal populations (**Fig. 5b** and **Extended Data Fig. 5b**). To increase pericyte specificity, we combined subtype-specific signatures with TAPS, restricting ECM- and IFN-associated activity scores to TAPS-positive perivascular cells (**Extended Data Fig. 5c,d**). This strategy markedly improved spatial resolution and confined both programs to discrete pericyte-associated niches (**Fig. 5c,d** and **Extended Data Fig. 5e,f**).

**Figure 5.**
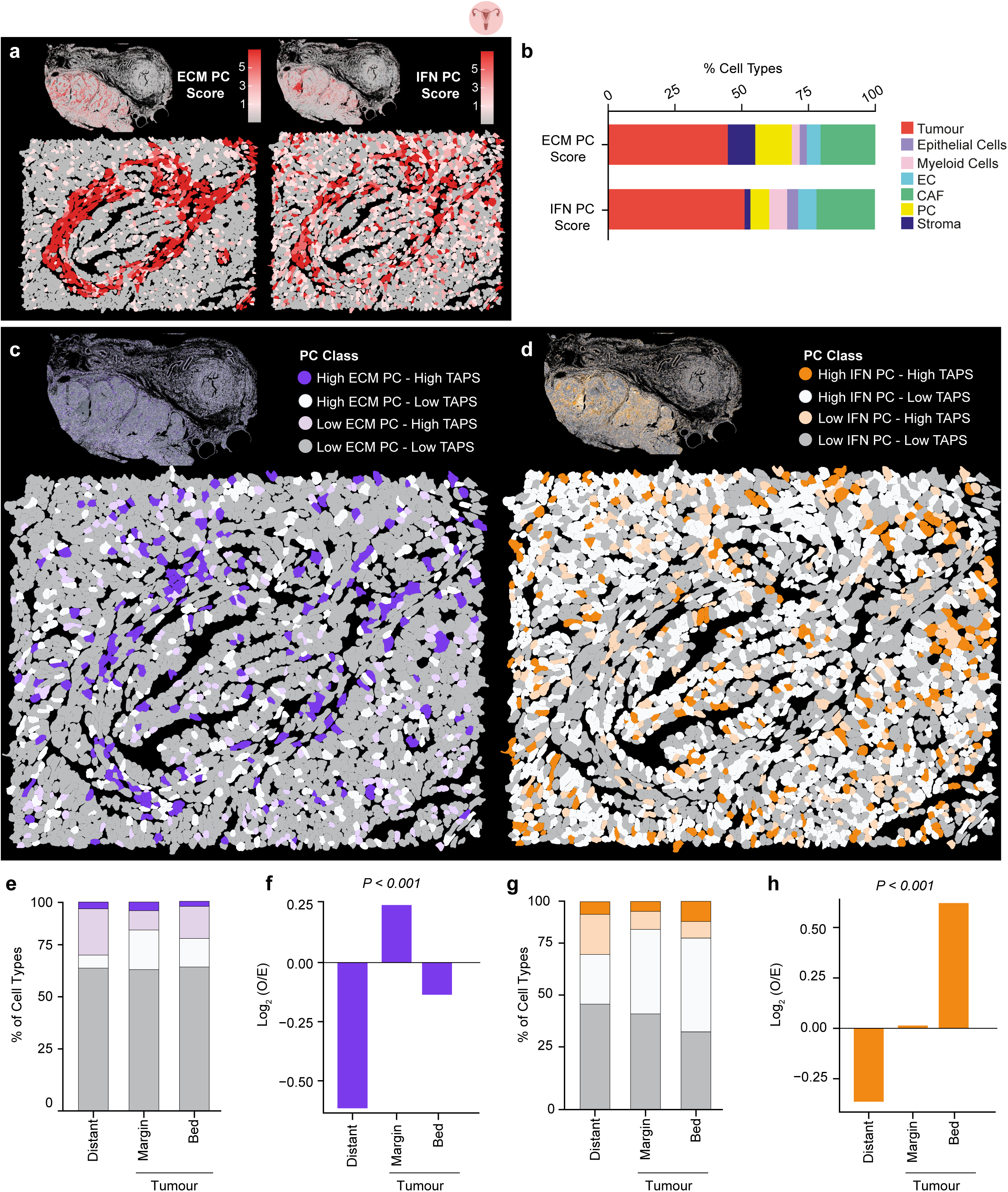
TAPS enables spatial resolution of specialised tumour-associated pericyte states. **a)** Spatial distribution of ECM (left) and IFN (right) pericyte (PC) signatures, quantified by UCell scores in the human ovary sample. Representative images showing the full tissue section (top) and a magnified inset (bottom) are provided for each signature. **b)** Stacked bar plot showing the classification of cell types exhibiting positive expression for ECM and IFN signatures. **c)** Spatial distribution of the ECM-associated PCs classified by the combined activity of the ECM and TAPS signatures. Representative images showing the full tissue section (top) and a magnified inset (bottom) are provided for each signature, in the human ovary sample. **d)** Spatial distribution of the IFN-responsive PC classified by the combined activity of the IFN and TAPS signatures. Representative images showing the full tissue section (top) and a magnified inset (bottom) are provided for each signature, in the human ovary sample. **e)** Stacked bar plot showing the distribution of ECM-associated PC across tumour zonation regions (distant, tumour margin and tumour bed) in the ovary sample. **f)** Bar plot showing the log_2_-enrichment score of ECM-associated PCs across the different tumour zones in ovary. Significance was determined by a chi-squared test (P<0.01). **g)** Stacked bar plot showing the distribution of IFN-responsive PCs across ovary tissue zones. **h)** Bar plot showing the log_2_-enrichment score of IFN-responsive PC across the different tumour zones in ovary. Significance was determined by a chi-squared test (P<0.01).

Having established the spatial localisation of pericyte states, we quantified their distribution across distinct tumour regions. ECM-associated pericytes displayed a consistent enrichment at the tumour margin across both ovarian and breast tumours, with additional enrichment in distant regions observed in breast tumours (**Fig. 5e,f** and **Extended Data Fig. 5g,h**, respectively). In contrast, IFN-responsive pericytes consistently accumulated within the tumour bed in both tumour types (**Fig. 5g,h** and **Extended Data Fig. 5i,j**). Together, these findings demonstrate that TAPS enables the spatial identification of specialised tumour-associated pericyte states *in situ* and reveals recurrent patterns of regional organisation across tumour architecture.

### Tumour-associated macrophages shape specialised pericyte states

We subsequently assessed the signalling mechanisms underlying the emergence of these specialised pericyte states in cancer by performing NicheNet^36^ ligand-receptor analysis using pericytes as receiver cells and all cellular compartments as senders (**Fig 6a**). Cross-tissue analysis of the top predicted ligands identified several recurrent signalling candidates, including *TGFB1*, *TNF*, *JAM2*, and members of the FGF family (**Fig 6b**). To prioritise signalling pathways most likely to influence the emergence of pericyte states, we next evaluated ligand-associated downstream target programs across tumour and non-tumour samples. *TGFB1* and *TNF* consistently displayed the strongest predicted ligand activities (**Fig. 6c**).

**Figure 6.**
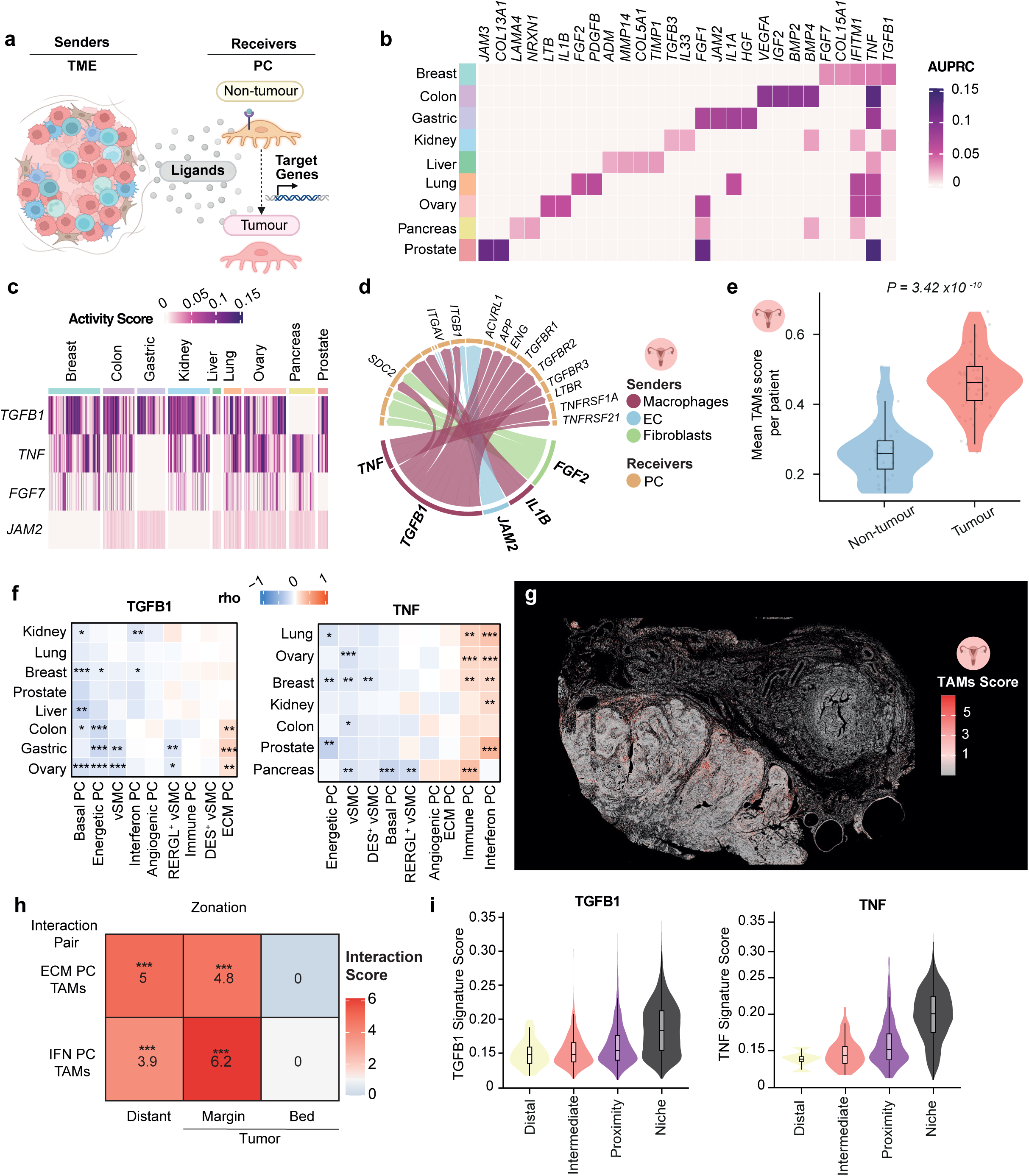
Tumour-associated macrophages shape specialised pericyte states. **a)** Schematic representation of NicheNet analysis, defining all cell types as senders and pericytes (PC) as receivers to assess transcriptomic changes post ligand-receptor binding. **b)** Heatmap showing the top 10 ligands across the nine atlas tissues; the purple gradient denotes AUPRC values. **c)** Heatmap of target genes for the most frequent ligands across tissues; the purple gradient represents gene activation levels upon ligand-receptor binding. **d)** Circle plot illustrating the top ligands and their corresponding receptors in ovary. **e)** Violin plot showing the mean UCell score per patient for the TAM signature within macrophage populations, comparing non-tumour and tumour samples. Statistical significance was determined using a Wilcoxon rank-sum test. **f)** Correlation heatmap showing the Spearman correlation between target gene scores for TGFB1 (right) and TNF (left) and the the log_2_FC of mural cell subtype markers. Statistical significance is indicated as: P ≤ 0.05 (*), P ≤ 0.01 (**), and P ≤ 0.001 (***). **g)** Spatial distribution of TAM signatures quantified by UCell scores; representative image shows the ovary tissue overview. **h)** Heatmap of interaction scores between ECM-associated and IFN-responsive PCs with TAMs across tissue zones in ovary. Statistical significance was assessed using a permutation test; FDR-adjusted P values are indicated as: adjusted P ≤ 0.05 (*), adjusted P ≤ 0.01 (), and adjusted P ≤ 0.001 (*). **i)** Violin plot with the average expression of *TGFB1* (left) and *TNF* (right) target gene signatures in the ovary tissue, categorized by distance between PC and TAMs: niche (<20 μm), proximity (20-50 μm), intermediate (50-100 μm), and distal (>100 μm).

Analysis of *TGFB1*- and *TNF*-associated ligand-receptor interactions revealed that macrophages were the predominant source of both ligands in most of the analysed tissues (**Fig. 6d, Extended Data Fig. 6a** and **Table S10**). To investigate the transcriptomic profile of macrophages under tumour conditions, we projected a curated tumour-associated macrophage (TAM) gene signature^37^ onto the macrophage compartment (**Table S2**). Tumour-derived macrophages exhibited significantly higher TAM signature activity than their non-tumour counterparts (**Fig. 6e** and **Extended Data Fig. 6b**). We next examined the expression of the predicted signalling ligands and found that macrophages expressed both *TGFB1* and *TNF* (**Extended Data Fig. 6c**), identifying these cells as a potential source of signalling contributing to pericyte reprogramming. To connect these interactions with mural cell states (**Fig. 4a**), we first confirmed preferential expression of *TGFB1*- and *TNF*-associated receptors within specialised mural cell subpopulations (**Extended Data Fig. 6d**). We then correlated ligand-target activity scores with the identity programs defining mural cell subpopulations. *TGFB1*-associated signalling preferentially correlated with ECM-associated pericytes (**Fig. 6f**, left), whereas *TNF*-associated signalling was most strongly associated with immune and IFN-responsive pericyte states (**Fig. 6f**, right). Together, these findings link TAM-derived signalling to specialised tumour-associated pericyte programs.

Projection of the TAM gene signature on the ovarian and breast spatial transcriptomic datasets revealed spatially restricted TAM-enriched regions within the myeloid compartment (**Fig. 6g** and **Extended Data Fig. 6e**). Spatial permutation analysis of nearest-neighbour distances confirmed significant enrichment of TAM-pericyte co-localisation at tumour-distant and margin regions across both tissues and specialised pericyte states, with no significant interactions detected within the tumour bed (**Fig. 6h** and **Extended Data Fig. 6f**). To further support a functional relationship between TAM proximity and pericyte state, we projected *TGFB1*- and *TNF*-associated target programs onto spatial transcriptomic data. Pericytes located near TAMs (<20 μm) exhibited the highest signature activity, which progressively declined with increasing distance from TAMs (**Fig. 6i** and **Extended Data Fig. 6g**). Together, these findings provide spatial support for potential TAM-pericyte interactions associated with specialised pericyte states.

### Distinct tumour ecosystems are associated with specialised pericyte states

To address whether specialised pericyte states were associated with distinct tumour ecosystems, spatial annotations were refined using a combination of pan-cancer single-cell identity programs and canonical marker gene expression (**Fig. 7a**, **Extended Data Fig. 7a,b** and **Table S2**). We interrogated whether ECM-associated and IFN-responsive pericytes occupied distinct spatial niches by quantifying the cellular composition within a 50 μm radius surrounding each pericyte (**Fig. 7b**). IFN-responsive pericytes were enriched within tumour cell-rich communities whereas ECM-associated pericytes exhibit a differential cancer-associated fibroblast (CAF)-rich neighbourhood across both tumour types (**Fig. 7c** and **Extended Data Fig. 7c**). Cumulative distance analyses confirmed the preferential association of ECM-associated pericytes with CAF-rich regions, whereas IFN-responsive pericytes were more closely associated with tumour-cell regions (**Fig. 7d** and **Extended Data Fig. 7d**). Spatial attraction-repulsion analyses further revealed a significant mutual exclusion between ECM-associated and IFN-responsive pericyte niches, indicating that these specialised populations occupy spatially segregated microenvironmental territories (**Fig. 7e** and **Extended Data Fig. 7e**).

**Figure 7.**
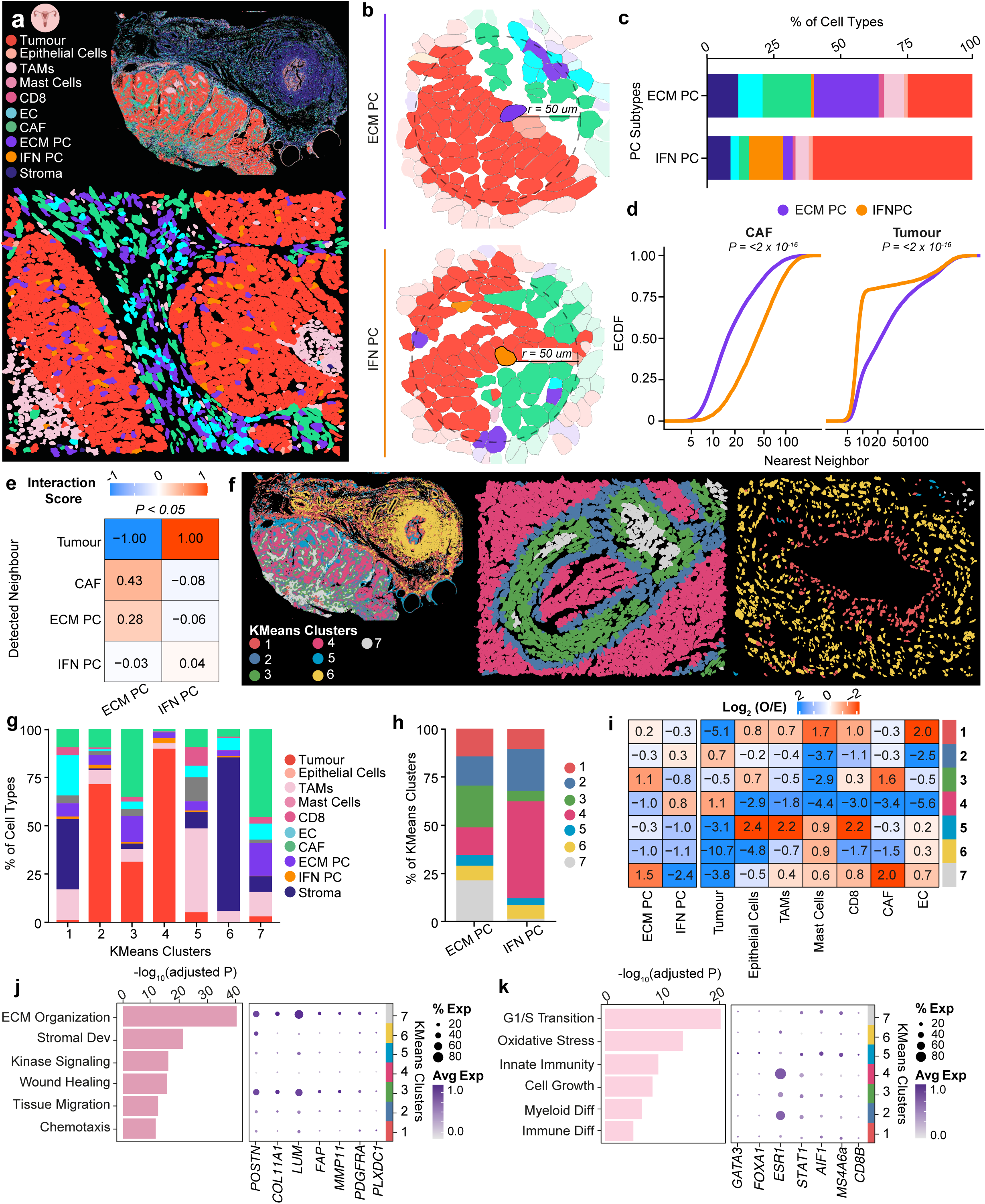
Distinct tumour ecosystems are associated with specialised pericyte states. **a)** Spatial distribution of the refined TME populations. Representative images showing the full tissue section (top) and a magnified inset (bottom) are provided in ovary. **b)** Zoom plot of the radius (r) representation, showing the spatial relationship between ECM-associated and IFN-responsive pericytes (PC) within a 50 µm radius, in ovary. **c)** Stacked bar plot showing the distribution of TME populations surrounding ECM-associated and IFN-responsive pericytes in ovary. **d)** Empirical cumulative distribution function (ECDF) curves showing the distances between ECM-associated and IFN-responsive PC with respect to CAFs and tumour cells in ovary. Statistical significance was estimated using the Kolmogorov-Smirnov test. **e)** Heatmap of interaction scores between ECM and IFN PCs with respect to tumour cells, CAFs, and other PCs, in ovary. Statistical significance was estimated using a permutation-based test. **f)** K-means spatial clustering representation. Representative images showing the full tissue section (left) and two magnified insets (middle and right) displaying spatial clusters in ovary. **g)** Stacked bar plot showing the distribution of TME cell types within the different defined K-means clusters in ovary. **h)** Stacked bar plot showing the distribution of ECM-associated and IFN-responsive PCs across the K-means clusters in ovary. **i)** Heatmap of the enrichment of the different cell types within the defined K-means clusters in ovary. **j)** Bar plot of overrepresented GO terms in clusters enriched in ECM-associated PCs (clusters 7 and 3, left) and dot plot showing the average expression (colour intensity) and percentage of expression (dot size) of the top marker genes across the different clusters (right) in ovary samples. **k)** Bar plot of overrepresented GO terms in clusters enriched in IFN-responsive PCs (cluster 4) and dot plot showing the average expression (colour intensity) and percentage of expression (dot size) of the top marker genes across the different clusters (right) in ovary samples.

To characterise these ecosystems in greater detail, we applied k-means clustering to group local neighbourhoods with similar cellular compositions and transcriptional features. This analysis identified seven spatial neighbourhood classes representing distinct TME architectures across tissues (**Fig. 7f** and **Extended Data Fig. 7f**). ECM-associated and IFN-responsive pericytes showed strikingly different distributions across these neighbourhood classes, with minimal overlap between the two populations (**Fig. 7g,h** and **Extended Data Fig. 7g,h**). ECM-associated pericytes preferentially localised to CAF-rich ecosystems characterised by ECM organisation, collagen remodelling, and fibrotic stromal programs, together with elevated expression of *POSTN*, *COL11A1*, *LUM*, *FAP*, *MMP11* and *PDGFRA* (**Fig. 7i,j** and **Extended Data Fig. 7i,j**). In contrast, IFN-responsive pericytes were enriched in tumour-cell ecosystems (**Fig. 7i** and **Extended Data Fig. 7i**), displaying inflammatory and immune-related programs, accompanied by expression of *STAT1*, *AIF1*, *MS4A6A* and *CD8B*. These ecosystems also retained tumour epithelial features marked by *GATA3*, *FOXA1* and *ESR1* expression (**Fig. 7k** and **Extended Data Fig. 7k**). Together, these findings indicate that specialised pericyte states are associated with distinct tumour ecosystems, linking ECM-associated pericytes to desmoplastic stromal environments and IFN-responsive pericytes to inflammatory tumour-immune niches.

### Specialised pericyte programs are associated with TME states and clinical outcome

We next investigated whether the specialised pericyte ecosystems identified by spatial transcriptomics could be detected in independent patient cohorts. To this end, we integrated ECM-associated and IFN-responsive pericyte scores with TME subtypes previously defined across >10,000 cancer patients^38^, and applied these classifications to TCGA bulk RNA-seq datasets spanning 24 cancer types (**Table S1**). In ovarian tumours, high ECM-associated pericyte activity was enriched in the fibrotic TME subtype, whereas high IFN-responsive pericyte activity was enriched in the immune and immune/fibrotic subtype (**Fig. 8a**). This pattern was consistent across all TCGA cancer types (**Fig. 8b**, and **Table S11)**, demonstrating that specialised pericyte states leave a transcriptional imprint detectable in bulk RNA-seq data. A pan-cancer meta-analysis confirmed the robustness of these findings (**Fig. 8c** and **Table S12**), indicating that the spatially identified ecosystem architecture is reproducible across independent patient cohorts.

**Figure 8.**
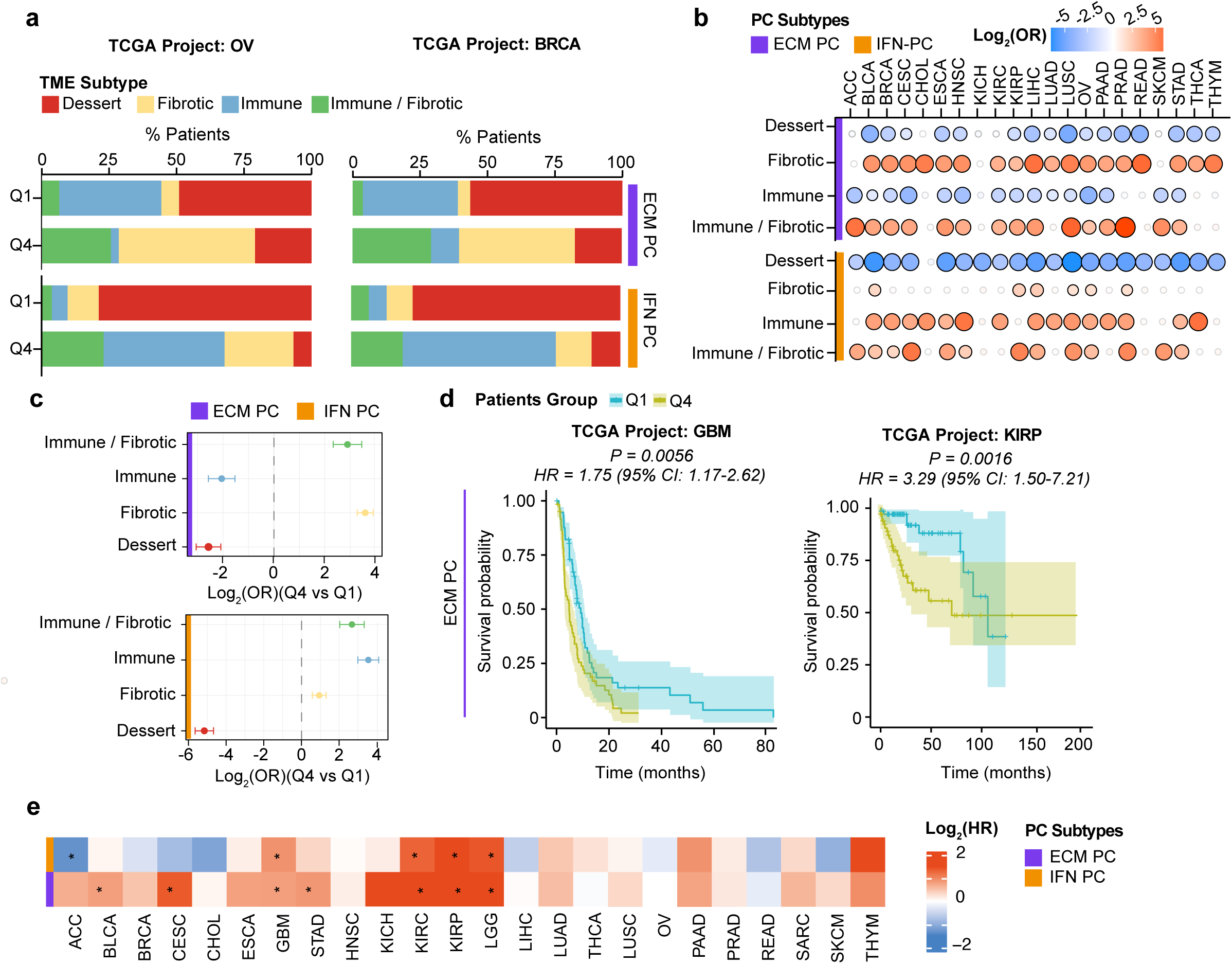
Specialised pericyte programs are associated with TME states and clinical outcomes. **a)** Stacked bar plot showing the distribution of patients stratified by high and low ECM-associated (top) and IFN-responsive (bottom) pericyte (PC) expression across different TME described by *Bagaev et al.* in the TCGA ovarian cancer (OV) cohort. **b)** Dot plot of the log2 odds ratio (OR) derived from Fisher’s exact test comparing the enrichment of TME subtypes between low- (Q1) and high-activity (Q4) PCs signature groups across TCGA cohorts. Significant associations (P < 0.05) are shown in red (positive association) or blue (negative association), whereas non-significant associations are shown in grey. **c)** Forest plot showing the enrichment of the different TME subtypes in ECM-associated (top) and IFN-responsive (bottom) pericyte stratifications. **d)** Kaplan-Meier estimates of disease-free survival in TCGA glioblastoma (GBM) cohort (GBM) and TCGA kidney renal papillary cell carcinoma (KIRP) cohort. Patients were stratified by ECM pericyte expression levels. Significance was determined by the Log-rank test. DFS is defined as the time from initial treatment to tumour recurrence or death from any cause. **e)** Heatmap of hazard ratios (HRs) for ECM-associated and IFN-responsive PC signatures across different TCGA cohorts. Statistical significance was assessed using a Cox proportional hazards model. Asterisks denote P < 0.05.

High ECM-associated pericyte scores were associated with reduced overall survival (**Fig. 8d**). Pan-cancer Cox proportional-hazard analyses confirmed this pattern, showing predominantly positive hazard ratios across both the tissue types represented in our atlas and additional cancers from independent TCGA cohorts, supporting a broadly adverse prognostic association of the ECM-associated pericyte program (**Fig. 8e**). In contrast, IFN-responsive pericyte programs showed no consistent association with patient outcome, displaying favourable, neutral or adverse prognostic effects depending on tumour type (**Fig. 8e**). Collectively, these results establish that specialised pericyte ecosystems are reproducible across independent patient cohorts and are associated with distinct TME states and clinical outcomes. Whereas ECM-associated pericytes represent a conserved adverse stromal programme, IFN-responsive pericytes display context-dependent associations with tumour immunity and prognosis.

## Discussion

The diverse developmental origins of pericytes^39^ and the absence of universal markers for their identification^9,12,13^ have complicated efforts to determine whether tumour-associated pericytes preserve their physiological tissue-specific identities or instead converge towards shared adaptive states. To address this question, we generated a dedicated pan-cancer atlas of the PVN spanning nine tissues, 911 specimens and 527,857 cells. Our atlas provides a comprehensive framework for understanding pericyte identity, reprogramming and specialised states in cancer. Among PVN populations, tumour-associated pericytes exhibited the strongest trend towards transcriptional convergence across tissues despite their marked heterogeneity in physiological settings ^11,14,15^. Pericytes exhibited substantially greater transcriptional diversity than vSMCs, aligning with recent pan-cancer studies that report extensive pericyte heterogeneity across tumour types^27,28^. Notably, mural cells remain systematically underrepresented in single-cell datasets owing to their low abundance and inefficient recovery during tissue dissociation. Indeed, nearly four million cells were required to recover approximately 50,000 pericytes in our atlas, underscoring both the technical challenges of studying these populations and the value of dedicated resources for their systematic investigation.

A major finding of our analysis is the identification of TAPS as a convergent transcriptional program shared across tumour-associated pericytes. Although several TAPS genes display tissue-specific expression under physiological conditions, tumour-associated pericytes exhibited markedly greater transcriptional concordance across tissues. One interpretation is that oncogenic cues coordinately activate pre-existing pericyte programs, enabling convergence towards a common adaptive state. Alternatively, TME-derived signals may independently drive pericytes towards similar transcriptional states across tissues. By integrating multiple genes into a coordinated signature, TAPS robustly distinguished pericytes from other PVN populations across independent single-cell and spatial transcriptomic datasets and outperformed canonical marker-based approaches in tumour contexts. Immunofluorescence validation confirmed expression of selected TAPS proteins in mural cells *in situ* and identified PLXDC1 as a tumour-enriched marker. However, variability across samples highlights the limitations of individual markers in identifying tumour-associated pericytes and underscores the value of combinatorial signatures, such as TAPS, for both defining these populations and prioritising new candidate markers.

We found that pericytes are recurrently expanded in tumours. This observation may appear surprising, given that reduced pericyte coverage has long been considered a hallmark of tumour vasculature^18,40^. However, diminished vascular coverage and reduced pericyte abundance are not necessarily equivalent phenomena. Our analyses suggest that at least part of the apparent reduction reported previously may reflect limitations of the canonical markers traditionally used to identify pericytes *in situ*, rather than a true loss of these populations. Indeed, recent work has shown that expression of some classical pericyte markers is diminished in tumours, whereas others are retained^16,41,42^, supporting the notion that reliance on individual markers can lead to misleading conclusions regarding pericyte abundance and function. Our finding that TAPS performs better in the TME than canonical pericyte markers indicates that tumour-associated pericytes adopt specialised transcriptional states that are only partially captured by classical markers, potentially contributing to their underestimation in previous studies.

Among the specialised pericyte states identified, ECM-associated and IFN-responsive pericytes emerged as convergent tumour-associated adaptations recurrently observed across cancers. ECM-associated pericytes displayed matrix-remodelling and fibrotic programs, consistent with previous reports describing stromal and fibrogenic pericyte phenotypes^12,19^. This population is also the most widespread across tumour types, suggesting that ECM remodelling represents a fundamental feature of tumour-associated pericyte adaptation rather than a tissue-specific response. Similar phenotypic transitions have been reported in fibrotic disorders, in which pericytes acquire myofibroblast-like features and contribute to ECM deposition^43,44^. Whether tumour-associated pericytes make a quantitatively important contribution to ECM remodelling relative to CAFs remains an open question. An equally intriguing issue is whether ECM-associated pericytes and CAFs represent distinct populations or share lineage relationships. Given their phenotypic similarities and close spatial association, lineage-tracing approaches will be required to define their respective contributions to ECM remodelling and their potential developmental relationships. The IFN-responsive pericytes exhibited inflammatory and interferon-associated programs. While this is consistent with previous studies showing that cancer-associated pericytes can regulate immune responses^12,45^, our analyses identify this activity as a recurrent IFN-responsive pericyte state shared across multiple tumour types.

Spatial analyses further showed that specialised pericyte states were embedded within distinct tumour ecosystems. ECM-associated pericytes preferentially localised to CAF-rich and desmoplastic regions, whereas IFN-responsive pericytes were enriched within inflammatory tumour environments. Their spatial partitioning suggests that specialised pericyte states are associated with distinct multicellular ecosystems rather than representing interchangeable transcriptional programs.

Predicted *TGFB1* and *TNF* signalling from TAMs further suggests that local microenvironmental cues contribute to the emergence of these specialised states. Consistent with recent spatial transcriptomic and multi-omics studies describing conserved multicellular ecosystems and stromal neighbourhoods across cancers^3,46^, these findings support a model in which tumour-associated pericytes integrate into specialised tumour ecosystems shaped by reciprocal interactions with neighbouring cells. The distinct correspondence of ECM-associated and IFN-responsive pericyte programs with TME states across independent patient cohorts further supports the biological and clinical relevance of these specialised ecosystems. In particular, the adverse prognostic impact of ECM-associated pericyte programs across multiple cohorts and cancer types demonstrates that stromal remodelling by pericyte subtypes is a clinically relevant feature of tumour progression.

Several limitations of this study warrant consideration. First, low TAPS activity was also observed in vSMCs and fibroblasts. While this overlap likely reflects the close transcriptional relationship among perivascular mesenchymal populations, it also underscores the importance of interpreting TAPS as a quantitative measure rather than a binary classifier. Second, spatial analyses were restricted to ovarian and breast tumours, extending this framework to additional samples and cancer types will be important for establishing the broader generalisability of the spatial organisation described here. Nevertheless, the convergence of single-cell, spatial, and bulk transcriptomic analyses across independent datasets supports the specialised pericyte states identified. Finally, although the association of these states with clinical outcome supports their biological relevance, direct experimental dissection of their causal contribution to tumour progression remains an important goal for future studies.

In summary, our study provides a comprehensive pan-cancer framework for understanding pericyte biology across human cancers. Despite their pronounced physiological heterogeneity, tumour-associated pericytes exhibited striking transcriptional convergence, revealing recurrent adaptive programs that transcend tissue boundaries. This convergence was accompanied by diversification into specialised states, including ECM-associated and IFN-responsive pericytes, which were embedded within distinct tumour ecosystems. The integration of single-cell, spatial, and ecosystem-level analyses further suggests that these specialised states are shaped by local microenvironmental signals and recurrent interactions with TAMs.

## Material and Methods

### Single-cell atlas construction

#### Single-cell RNA-seq data acquisition and preprocessing

Publicly available scRNA-seq datasets generated using droplet-based technologies (10X Genomics, Seq-Well, BD Rhapsody, GEXSCOPETM, and inDrops) were retrieved from public repositories, including Gene Expression Omnibus, CellxGene, Zenodo, and the Human Cell Atlas. The studies included in the atlas are summarised in **Table S1**. Patient-level information, including sample annotations and clinical metadata, is provided in **Table S1**. All analyses were performed using Seurat (v4.4.0). For each dataset, Seurat objects were generated, and metadata were harmonised to ensure compatibility across studies. Quality-control filtering was performed independently for each dataset following current single-cell transcriptomic best-practice recommendations^47^. Cells with low transcript recovery (<250 UMIs), high mitochondrial content (>20%), or abnormal transcriptomic complexity (complexity index >0.8) were excluded. The complexity index was calculated as previously described^48^ as the ratio of the log_10_-transformed number of UMIs to the number of detected genes and was used to identify low-quality outlier cells. Genes detected in fewer than five cells were removed prior to downstream analyses. To ensure sufficient representation of rare stromal populations, patients contributing fewer than 1,000 cells were excluded. Potential doublets were identified independently for each patient using DoubletFinder (v2.0.3). Doublet frequencies were evaluated across tissues and stromal populations to assess potential technical biases. Gene expression values were normalised using Seurat’s *LogNormalize* method with a scale factor of 10,000. Highly variable genes were identified using the *FindVariableFeatures* function with the variance-stabilising transformation method. Normalised data were scaled and subjected to principal component analysis (PCA). The optimal number of principal components was estimated independently for each dataset using variance-explained criteria and generally ranged from 15 to 25 components. Cell clustering was performed using the Louvain algorithm. Nearest-neighbour graph was constructed from the PCA embeddings. Clustering was evaluated across multiple resolutions (0.1-0.7) and the resolution providing optimal separation of the major TME compartments was selected for downstream analyses. Uniform Manifold Approximation and Projection (UMAP) was subsequently used to visualize the resulting cellular landscape. Following quality control and preprocessing, datasets belonging to the same tissue were merged into a unified Seurat object to generate tissue-specific atlases. To minimize technical variation arising from differences between patients, batch correction was performed using Harmony with patient identity specified as the batch variable^49^. The Harmony-corrected embeddings were subsequently used for graph construction, clustering and UMAP visualization. Major TME compartments were subsequently annotated based on canonical marker gene expression (**Table S2 and S3**). The resulting tissue-specific atlases provided the framework for downstream PVN extraction, cross-tissue integration and subsequent analyses.

#### Perivascular niche extraction

To specifically investigate the PVN, ECs, fibroblasts and mural cells were extracted from each tissue-specific atlas using Seurat’s *subset* function. The resulting tissue-specific PVN datasets were reprocessed independently following the same analytical workflow described above, including identification of highly variable genes, scaling, principal component analysis, Harmony batch correction, and graph-based clustering. Major perivascular populations were subsequently annotated based on canonical marker gene expression (**Table S2**).

#### Cross-tissue integration of the perivascular niche

Following tissue-specific annotation, all PVN datasets were merged into a unified cross-tissue dataset. The merged object was reprocessed using the same analytical workflow, including highly variable gene selection, scaling, principal component analysis, Harmony batch correction and graph-based clustering.

#### Cell population annotation and marker identification

To define the molecular identity of cellular populations, cluster-specific marker genes were identified using Seurat’s *FindMarkers* and *FindAllMarkers* functions. Differential gene expression (DEG) was assessed using the Wilcoxon rank-sum test with Bonferroni correction for multiple testing. Marker genes were identified using a one-versus-all comparison strategy, in which each population was compared against all remaining cells in the dataset.

#### Differential abundance analysis

Differences in cell population abundance were quantified using the propeller method implemented in the speckle package^29^. Cell type proportions were estimated at the sample level and compared between conditions using an empirical Bayes moderated t-test. Statistical significance was determined using adjusted p-values (<0.05).

### Pericyte identity program analysis

#### Cell type-specific differential expression analysis

To define the transcriptional identity of PVN populations, gene-expression analyses were performed using a one-versus-all strategy. Within each tissue and condition (tumour or non-tumour), each cell population (ECs, fibroblasts, pericytes and vSMC) was compared against the cells from all other perivascular populations from the same tissue and condition. DEG was performed using Seurat’s *FindAllMarkers* function (Wilcoxon rank-sum test with Bonferroni correction). Marker genes were identified using a one-versus-all comparison strategy, in which each population was compared against all remaining cells in the dataset. For each population and condition, significantly enriched genes (adjusted P < 0.05) were ranked by log_2_ fold change (log₂FC) and used in downstream analyses.

#### Cross-tissue identity program analysis

Gene sets identified in each population were subsequently compared across tissues to identify common and tissue-specific transcriptional programs. For each population and condition separately, the top 250 significantly upregulated genes ranked by log₂FC were retained, and intersections across tissues were quantified using UpSet plots. This threshold corresponded approximately to the upper 10% of DEG across tissues and was selected to ensure consistent gene-set sizes for cross-tissue conservation analyses. Common genes identified across tumour pericytes were used to define the TAPS, which was subsequently used for downstream analyses. Common gene programs derived for ECs, fibroblasts and vSMC are reported in **Table S5**.

#### Downsampling-based robustness analysis

To assess whether differences in cell numbers between non-tumour and tumour conditions affected the identification of common transcriptional programs, a downsampling analysis was performed. For each tissue, tumour pericytes were randomly subsampled to match the number of non-tumour pericytes. This procedure was repeated 1,000 times. For each iteration, gene-expression analysis was performed as described above and the top 250 significantly upregulated genes were retained. Cross-tissue intersections were subsequently recalculated to quantify the number of common genes identified across tumour tissues. The distribution of common genes obtained across iterations was compared with the number observed in the original analysis. An enrichment score was calculated as the ratio between the mean number of common genes identified across downsampling iterations and the number observed in the reference condition. Empirical p-values were estimated as the fraction of iterations yielding fewer common genes than observed in the reference condition.

#### Human samples for immunofluorescence

Formalin-fixed paraffin-embedded (FFPE) human breast tumour samples were obtained from Germans Trias i Pujol Hospital and General University Hospital of Albacete following approval by the corresponding institutional ethics committees. In addition, commercially available human tissue microarrays (TMA) were purchased from TissueArray.Com (Derwood, MD, USA). Sample information can be found in **Table S1**. Both breast and ovarian (TMA and FFPE) human tissue samples were processed for immunofluorescence staining as follows. Tissue sections were baked at 60°C for 30 min to 1 hour, deparaffinized in xylene, and rehydrated through a graded ethanol series to distilled water. Antigen retrieval was performed in preheated 10 mM citrate buffer, pH 6.0, using a pressure cooker for 10 min, followed by gradual cooling for 15 min. Slides were washed in PBS, permeabilized and blocked for 1 h at room temperature in PBS containing 0.3% Triton X-100 and 5% BSA, and incubated overnight at 4°C with primary antibodies diluted in the same solution. After three 5-min washes in PBS-T, slides were incubated with secondary antibodies for 2 h at room temperature, washed again in PBS-T, rinsed in PBS, mounted in Fluoromount-G (0100-01, Southern Biotech) and stored at 4°C for at least 24 h before imaging. Microscopic analysis was performed using a Zeiss LSM900 confocal microscope, where high-resolution images were acquired using a 20× objective. Between four and seven blood vessels per patient were included in the analysis, depending on vessel availability within each section, with a minimum of three patient samples analysed for each antibody panel. Images were processed and quantified using Imaris 10.1 software. Mean fluorescence intensity (MFI) of the indicated marker was quantified within the αSMA or NOTCH3 surface in co-localization with CD31+ cells. Data analysis was performed using GraphPad Prism (v10.4.1 for Windows). Nuclei were stained with 4,6-diamidino-2-phenylindole (DAPI). The antibodies used were the following. Primary antibodies: CD31/PECAM-1 (1:100; AF3628, R&D Systems); Cy3-conjugated αSMA (1:200; C6198, Sigma-Aldrich); TEM7/PLXDC1 (1:100; MA1-91725, Invitrogen); CDH6 (1:100; AF2715-SP, R&D Systems); ITGA7 (1:100; NBP1-86118, Novus Biologicals); FOXS1 (1:100; 16234-1, Proteintech). Secondary antibodies: Donkey anti-goat Alexa Fluor 488 (1:200; A11055, Invitrogen); donkey anti-rabbit Alexa Fluor 488 (1:200; A11055, Invitrogen); donkey anti-rabbit Alexa Fluor 647 (1:200, A32795, Invitrogen); donkey anti-sheep Alexa Fluor 647 (1:200; 10729004, Abcam).

#### Gene signature scoring

To quantify the activity of predefined transcriptional programs at the single-cell level, gene signature scoring was performed using UCell^50^, a rank-based method for single-cell gene set enrichment analysis.

#### TAPS activity analysis

To evaluate TAPS activity across biological conditions and mural cell populations, single-cell expression scores were first computed using the same methodology described above for general gene signature scoring. Subsequently, individual patient-level mean TAPS scores were calculated independently for each cell population. These patient-level mean scores were used as biological replicates for all subsequent statistical analyses to avoid pseudoreplication arising from cell-level comparisons. Statistical significance was assessed using the Wilcoxon rank-sum test. TAPS activity was first compared across mural cell populations, including pericytes, endothelial cells, fibroblasts and vSMC, to evaluate its cell type specificity. Subsequently, differential TAPS activity between tumour and non-tumour conditions was assessed specifically within the pericyte population. Both analyses were performed across the entire pan-cancer dataset and independently within each tissue to evaluate the consistency of the observed patterns across anatomical sites.

#### Logistic regression classifier analysis

To evaluate the ability of TAPS to identify pericytes, supervised logistic regression models were constructed independently in non-tumour and tumour samples. Cells from all tissues were classified as either pericytes or non-pericyte perivascular populations (ECs, fibroblasts and vSMC), while UCell-derived signature scores were used as predictor variables. For each analysis, cells were randomly divided into training (80%) and testing (20%) datasets using stratified sampling to preserve class proportions. Logistic regression models were fitted on the training set and evaluated on the testing set. Model performance was assessed using precision-recall (PR) curves and the corresponding area under the precision-recall curve (AUPRC), a metric well-suited for imbalanced classification settings given the minority representation of pericytes within PVN populations. The same classification framework was subsequently applied to the independent validation cohort to assess the reproducibility and generalizability of TAPS-based pericyte identification across datasets. To benchmark the discriminatory capacity of TAPS, a canonical pericyte marker signature (*ABCC9*, *ANPEP*, *CD248*, *CSPG4*, *HIGD1B*, *KCNJ8*, *NDUFA4L2*, *PDGFRB*, *RGS5* and *NOTCH3*) was quantified using UCell in the independent validation cohort. TAPS and canonical marker signature scores were compared using precision-recall curves and AUPRC to evaluate their relative ability to discriminate pericytes from other perivascular cell populations under both non-tumour and tumour conditions.

#### Independent dataset validation

To validate the robustness of TAPS in an independent setting, we analysed the glioblastoma scRNA-seq dataset^34^ (GSE256493). Processed data and cell type annotations generated by the original study were used directly. This dataset was subsequently used to validate TAPS activity, evaluate logistic regression classifier performance and benchmark TAPS against canonical pericyte marker signatures. To evaluate TAPS activity across conditions, we first quantified the mean TAPS score at the patient level, thereby treating each patient as the independent biological replicate. We additionally examined cell-level distributions to assess whether the same trend was consistently observed across individual cells. Statistical significance was assessed using the Wilcoxon rank-sum test.

### Mural cell heterogeneity analysis

#### Mural cell subsetting and annotation

To investigate heterogeneity within the mural compartment, pericytes and vSMC were extracted from the integrated PVN dataset. The resulting dataset was reprocessed using the analytical workflow described above, including highly variable gene selection, scaling, Harmony batch correction, dimensionality reduction and graph-based clustering. Clusters were initially annotated as pericytes or vSMC based on established canonical marker genes (**Table S2**). To further characterise mural cell heterogeneity, cluster-specific marker genes were identified using the DEG framework described above and subjected to functional enrichment analysis. The combined interpretation of canonical markers, transcriptional signatures and enriched biological processes was used to assign biological identities to the resulting mural cell populations.

#### Functional enrichment analysis

Functional enrichment analyses were performed using the enrichGO function from the clusterProfiler package^51^ using Gene Ontology Biological Processes (GO-BP) as the reference ontology (parameters: ont = "BP", OrgDb = org.Hs.eg.db, keyType = "SYMBOL", minGSSize = 10 and maxGSSize = 250). Terms with adjusted P < 0.05 were considered significant.

#### Cross-study comparison of mural cell states

To compare mural cell populations identified in this study with previously reported mural cell states, marker genes previously described^27,28^ were retrieved and their corresponding log₂FC values were extracted. For each mural cell population identified in our dataset, log₂FC values for the same genes were calculated using the DEG framework described above. Similarity between populations was assessed using pairwise Spearman correlation analyses.

#### Transcription factor regulon inference and activity analysis

To investigate the transcriptional regulatory programs underlying mural cell heterogeneity, we performed SCENIC^35^ analysis on the integrated mural cell dataset using the pySCENIC package (v0.12.1). Initial gene regulatory networks were inferred using GRNBoost2, followed by a refinement step to identify direct-binding targets through cisTarget motif enrichment analysis against human regulatory databases. Regulon activity was subsequently quantified at the single-cell level using AUCell. Finally, regulon activity scores were aggregated by mural cell population.

### Spatial transcriptomic analysis

#### Spatial transcriptomic data processing

Spatial transcriptomic data were obtained from 10x Genomics Xenium 5K datasets from breast and ovarian tumours and analysed using Seurat (v5.0.0) (**Table S1**). Cells with fewer than 20 detected transcripts were excluded. Data were log-normalized, highly variable genes were identified, and dimensionality reduction was performed using PCA. To facilitate analysis of the large-scale Xenium datasets, sketch-based subsampling implemented in Seurat was used to generate representative low-dimensional embeddings. Graph-based clustering was performed across multiple resolutions and visualized using UMAP embeddings and tissue coordinates. To resolve transcriptionally distinct cellular states within specific tissue niches, selected populations were subjected to iterative subclustering, including re-identification of highly variable genes, dimensionality reduction, clustering and marker gene analysis. Cell identities were assigned using the same canonical marker genes employed for scRNA-seq annotation (**Table S2**). Refined annotations obtained from these subset analyses were subsequently projected back onto the full Xenium dataset, allowing visualisation and quantification of the identified populations within the complete spatial tissue context. For visualisation purposes, selected regions of interest were extracted from the full Xenium dataset and displayed using their original spatial coordinates to provide higher-resolution views of local cellular organisation (**Table S13**).

#### Spatial tissue zonation

To characterise the spatial organisation of the TME, a distance-based zonation framework was implemented. Tumour cells were identified using a combination of tumour-specific transcriptional signatures (**Table S2**). Spatially connected tumour regions were subsequently defined by constructing neighbourhood graphs of tumour cells and identifying connected tumour masses. For each cell in the dataset, the minimum Euclidean distance to the nearest tumour cell was calculated. Based on this distance, cells were assigned to one of three spatial compartments: tumour bed (tumour cells), tumour margin (non-tumour cells located within 100 μm of the tumour boundary), and distant (non-tumour cells located more than 100 μm from the tumour boundary).

#### TAPS-based spatial pericyte identification

To quantify the relative coverage of annotated pericytes by TAPS and canonical markers in spatial transcriptomic data, we used the published cell type annotations from the ovarian Xenium dataset as the reference, defining the annotated pericyte population as the benchmark for comparison. Because transcriptional signatures and individual marker genes exhibit continuous rather than binary activity, the upper quartile (75th percentile) of the corresponding UCell score or gene expression distribution was used to define cells with robust signature activity or marker expression (**Extended Data Fig. 3f**). Coverage was then calculated as the proportion of annotated pericytes classified as TAPS-positive, *PDGFRB*-positive or *NOTCH3*-positive.

#### Co-expression-based identification of specialised pericyte ecosystems

To identify ECM-associated and IFN-responsive pericyte ecosystems in spatial transcriptomic data, ECM and IFN transcriptional programs, derived from the single-cell atlas, were represented by the top 200 most significantly upregulated genes for each population. Signature activities were quantified as described above for gene signature scoring. Cells with robust activity for each signature were defined using the upper quartile (75th percentile) of the corresponding signature score distribution (**Extended Data Fig. 3f** and **Extended Data Fig. 5c,d**). Because scRNA-seq analyses demonstrated that ECM-associated and IFN-responsive pericytes represent transcriptionally distinct populations, mutually exclusive classification criteria were applied in the spatial datasets. Specifically, specialised ECM-associated pericytes were defined as cells with high TAPS and ECM activity together with low IFN activity, whereas specialised IFN-responsive pericytes were defined as cells with high TAPS and IFN activity together with low ECM activity. This strategy was implemented to preserve the mutually exclusive identities observed in the higher-resolution single-cell data while accounting for the lower transcriptional resolution of Xenium spatial transcriptomics.

#### Spatial enrichment of specialised pericyte states

To assess the spatial distribution of specialised pericyte states, the frequency of dual-positive cells was quantified across the previously defined tumour zonation compartments. For each zone, observed dual-positive cell counts were compared with expected counts derived from the global distribution of cells across the tissue using a chi-squared test of independence. Spatial enrichment was quantified as the log_2_ ratio between observed and expected cell counts.

### Spatial neighbourhood and interaction analysis

#### Spatial niche definition

To characterise the local microenvironments associated with specialised pericytes, spatial neighbourhoods were constructed using a 50 μm interaction radius. For each ECM-associated and IFN-responsive pericytes, the abundance of neighbouring cellular populations within this radius was quantified, generating a local neighbourhood composition profile. Pericytes sharing similar neighbourhood compositions were subsequently grouped using k-means clustering. The number of clusters (k = 7) was selected based on the observation that neighbourhood structures remained largely unchanged with further increases k increased, indicating convergence towards stable spatial patterns. To annotate the resulting spatial niches, cell type enrichment was quantified by comparing the observed abundance of each cellular population within each niche with the abundance expected from its global distribution across the tissue. Enrichment scores were calculated as the log_2_ ratio between observed and expected counts, with pseudocount correction. To characterise the biological identity of each spatial niche, marker genes were identified by gene expression analysis and subsequently subjected to GO enrichment analysis, as described in the Functional Enrichment Analysis section.

#### Spatial interaction analysis

Spatial interactions involving ECM and IFN-specialised pericytes were assessed using a radius-based neighbourhood approach implemented in scimap. Cell-cell interactions were evaluated within a 50 μm radius and compared against randomised spatial distributions generated through permutation testing. Interaction scores were used to quantify attraction or avoidance between ECM-associated pericytes, IFN-responsive pericytes and the surrounding cellular populations. Statistical significance was assessed using a permutation test (n = 1,000 permutations), with empirical P values calculated by comparing the observed interaction frequencies to the null distributions generated from randomised cell locations. Analyses were conducted globally and independently within each zonation compartment.

### Cell-cell communication analysis

#### Ligand-receptor interaction inference

To identify signalling pathways potentially associated with tumour-induced pericyte reprogramming, ligand-receptor analyses were performed using NicheNet^36^. For each tissue, pericytes were designated as the receiver population, while all remaining annotated cell types were considered potential sender populations. Prior to gene expression analysis, cells identified as doublets were excluded and not included in the definition of the receiver population. DEG between tumour and non-tumour pericytes were then used as the target gene set, and ligand activity was estimated using the NicheNet prior model, which integrates ligand-receptor interactions, intracellular signalling, and gene regulatory networks. Candidate ligands were ranked according to their predicted ability to explain the observed transcriptional changes in tumour-associated pericytes.

#### Definition of tumour ligand response programs

To investigate downstream transcriptional programs associated with candidate ligands, tumour ligand-response programs were defined using NicheNet-predicted target genes. For each ligand, the target genes identified across tissues were aggregated and ranked by their predicted regulatory potential.

#### Association of ligand-response programs with mural cell states

To investigate whether ligand-response programs were preferentially associated with specific mural cell states, predicted ligand target genes were compared with mural cell subtype marker genes identified in **Fig 4**. Within each tissue, the overlap between ligand-response programs and mural cell subtype signatures was determined, and Spearman correlation coefficients were calculated using the corresponding log₂FC values.

### Spatial validation of TAM-pericyte signalling

#### TAM characterization

TAMs were identified by projecting a curated TAM gene signature onto the annotated macrophage compartment, with signature activity quantified as described above for gene signature scoring. The signature comprised TAM-associated genes reported^37^, together with additional established macrophage/TAM markers, and is provided in **Table S2**. To compare TAM activity between tumour and non-tumour conditions, patient-level mean signature scores were calculated by averaging single-cell scores across macrophages from each patient.

These patient-level mean scores were used as biological replicates, and statistical significance was assessed using the Wilcoxon rank-sum test.

#### TAM proximity analysis

To characterise the spatial relationship between specialised pericytes and TAMs, Euclidean distances were calculated from each ECM-associated and IFN-responsive pericyte to the nearest TAM within each Xenium sample. Pericytes were stratified into four distance intervals based on their proximity to TAMs: niche (<20 μm), proximity (20-50 μm), intermediate (50-100 μm), and distal (>100 μm).

#### TAM-pericyte interaction analysis

To assess whether TAM-pericyte associations exceeded those expected by chance, spatial interaction analysis was performed independently within each tumour zonation compartment. A local spatial graph was first constructed for each tissue zone using a 50 μm interaction radius. Interactions between ECM-associated or IFN-responsive pericytes and TAMs were then evaluated using the *testInteractions* function from imcRtools, with statistical significance determined by 1,000 permutation tests.

### Clinical and translational analyses

#### Bulk transcriptomic signature scoring and patient stratification

To assess the clinical relevance of specialised mural cell states, ECM-associated and IFN-responsive pericytes scRNA-seq signatures were projected onto bulk RNA-seq from all available TCGA solid tumour types. Signature activities were quantified using single-sample Gene Set Enrichment Analysis (ssGSEA). Pericyte subtype signatures were defined using the top 200 most significantly upregulated marker genes identified for each subtype by Seurat’s *FindMarkers* function applied to the scRNA-seq dataset. For patient stratification, samples were ranked according to their ssGSEA enrichment scores. To maximise separation between groups while minimising ambiguity associated with intermediate values, only patients belonging to the lowest (Q1) and highest (Q4) quartiles were retained for downstream analyses.

#### Tumour microenvironment ecosystem association analysis

To investigate the association between specialised pericyte states and TME ecosystem composition across TCGA solid tumour cohorts, previously published molecular functional phenotype (MFP) classifications^38^. This MFP were integrated with the patient stratification framework defined above. ECM-associated and IFN-responsive pericyte signatures were analysed independently, and the distribution of MFP classes was compared between low- (Q1) and high-activity (Q4) patient groups using Fisher’s exact test.

#### Survival analysis

The prognostic significance of pericytes signatures was evaluated across TCGA solid tumour cohorts using disease-free survival as the clinical endpoint. Kaplan-Meier survival curves, log-rank tests and Cox proportional hazards regression models were implemented using the survival package to estimate survival differences, hazard ratios and corresponding 95% confidence intervals between groups.

## Data availability

All datasets analysed in this study are publicly available and summarised in **Table S1**, which includes accession numbers and data sources. Single-cell RNA sequencing datasets were obtained from publicly available repositories, while spatial transcriptomic datasets were downloaded from the 10x Genomics data portal. TCGA bulk RNA-seq data were obtained from the Genomic Data Commons portal.

## Code availability

All custom scripts used for data processing, integration, gene-expression analysis, spatial transcriptomic analyses, cell-cell communication inference, and downstream statistical analyses are available at GitHub: https://github.com/anemartinezlarrinaga2898/AnalysisByTissue. The repository contains the code required to reproduce all analyses presented in this study.

## Acknowledgements

We thank CERCA Program/Generalitat de Catalunya and the Josep Carreras Foundation for their institutional support. We thank the members of Graupera and Mendizabal laboratories for critical feedback. We thank the Biobank of the General University Hospital of Albacete for their technical support. Figures **1a**, **6a** and the tissue icons were created in BioRender. Graupera, M. (2026) https://BioRender.com/vn3uctf.

## Authors’ contributions

M.G., A.M-L., I.M., and S.Camargo. were the main contributors to the conception, design, acquisition, and interpretation of the data and to writing the article. A.M-L., S.Camargo., M.S., S. Cervilla., S. G-L, A.M-R, L.G., H.v.S. performed experiments and data analysis with input from M.G, I.M. E.P-P, V.R., A.C. gave scientific input. R.A.B-A., E.M. and E.M.G-M. provided human samples.

## Ethical compliance

This study was conducted in accordance with the Declaration of Helsinki and all applicable national regulations governing biomedical research involving human subjects. Clinical patient samples were collected with approval from the Ethical Committee of Germans Trias i Pujol Hospital (CEIC #PI-21-230) and General University Hospital of Albacete (CEIC #2020/06/071, #2022-087, and #2023-058).

Written informed consent was obtained from all participants prior to their inclusion in the study.

