## Supplementary Figures for "A pan-cancer single-cell atlas of pericytes"

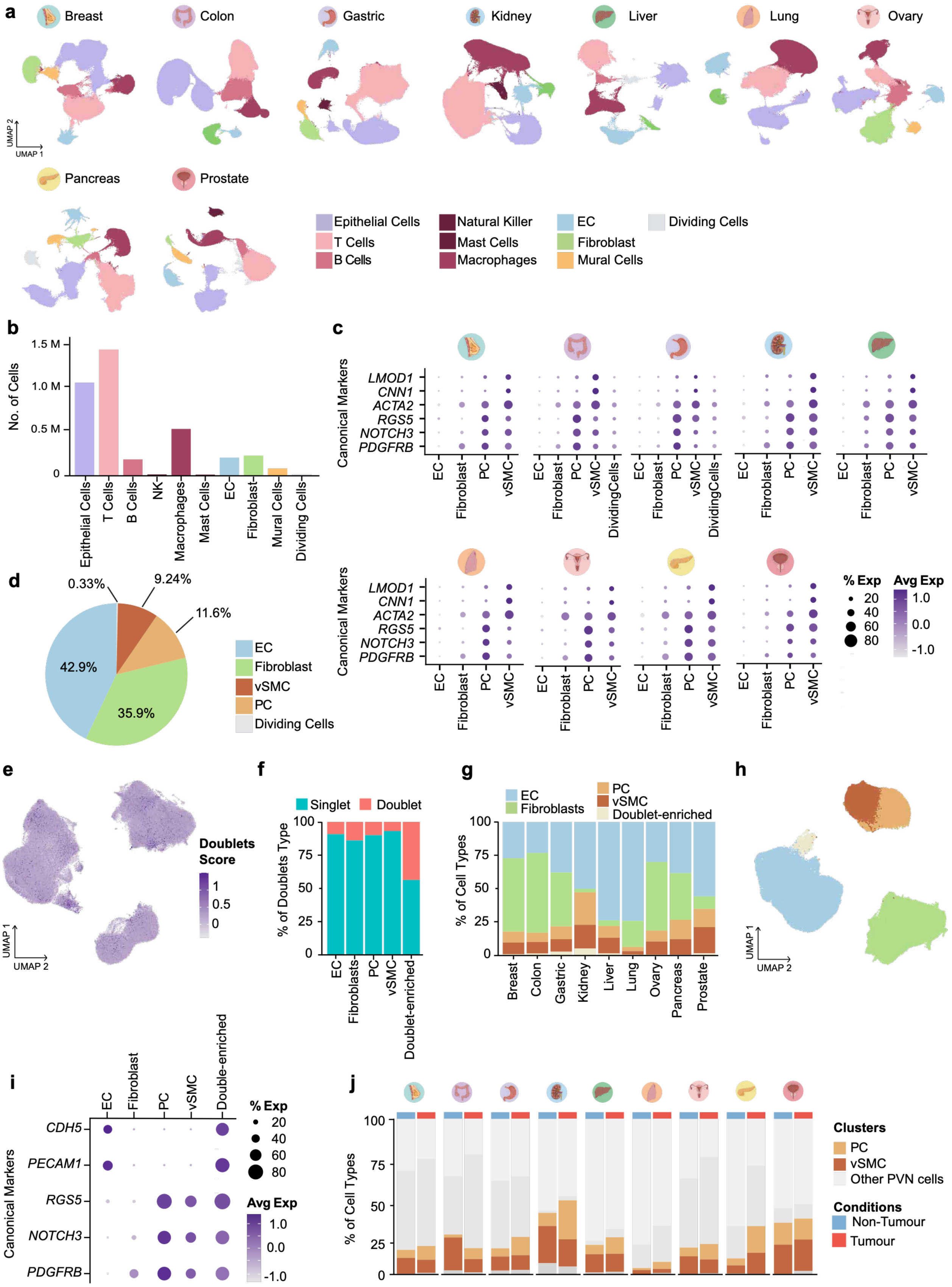
a
Breast
Colon
Gastric
Kidney
Liver
Lung
Ovary
Pancreas
Prostate
UMAP 1
UMAP 2
Epithelial Cells
T Cells
B Cells
Natural Killer
Mast Cells
Macrophages
EC
Fibroblast
Mural Cells
Dividing Cells
b
No. of Cells
1.5 M
1.0 M
0.5 M
0
Epithelial Cells
T Cells
B Cells
NK
Macrophages
Mast Cells
EC
Fibroblast
Mural Cells
Dividing Cells
c
Canonical Markers
LMOD1
CNN1
ACTA2
RGS5
NOTCH3
PDGFRB
EC
Fibroblast
PC
vSMC
DividingCells
% Exp
20
40
60
80
Avg Exp
1.0
0.0
-1.0
d
0.33%
9.24%
11.6%
42.9%
35.9%
EC
Fibroblast
vSMC
PC
Dividing Cells
e
Doublets Score
1
0.5
0
UMAP 1
UMAP 2
f
Singlet
Doublet
% of Doublets Type
100
75
50
25
0
EC
Fibroblasts
PC
vSMC
Doublet-enriched
g
EC
Fibroblasts
PC
vSMC
Doublet-enriched
% of Cell Types
100
75
50
25
0
Breast
Colon
Gastric
Kidney
Liver
Lung
Ovary
Pancreas
Prostate
h
UMAP 1
UMAP 2
i
EC
Fibroblast
PC
vSMC
Double-enriched
Canonical Markers
CDH5
PECAM1
RGS5
NOTCH3
PDGFRB
% Exp
20
40
60
80
Avg Exp
1.0
0.0
-1.0
j
% of Cell Types
100
75
50
25
0
Clusters
PC
vSMC
Other PVN cells
Conditions
Non-Tumour
Tumour

**Extended Data Fig. 1:** A pan-cancer atlas reveals selective expansion of pericytes.

**a)** UMAP plots showing the identification of the main TME compartments across all tissues, organized alphabetically: breast, colon, gastric, kidney, liver, lung, ovary, pancreas, and prostate. **b)** Bar plot representing the number of cells in each of the main annotated compartments across all tissues. **c)** Dot plot showing the average expression (colour scale) and percentage of expression (dot size) of canonical markers used to identify pericytes (PC; *RGS5*, *NOTCH3*, *PDGFRB*) and vSMC (*ACTA2*, *CNN1*, *LMOD1*). **d)** Pie chart representing the proportions of the main cell subtypes within the unified PVN dataset. **e)** UMAP plot displaying the DoubletFinder score in the integrated PVN dataset. **f)** Bar plot showing the percentage of singlets and doublets identified in the different populations of the unified PVN atlas. **g)** Bar plot showing the distribution of the identified populations in the unified PVN dataset across all analysed tissues. **h**) UMAP plot showing the distribution of the main PVN compartments after doublet removal. **i)** Dot plot showing the expression of EC markers (*PECAM1*, *CDH5*) and PC markers (*RGS5*, *NOTCH3*, *PDGFRB*). Dot size represents the percentage of cells expressing each marker, while colour intensity indicates the average expression level. **j)** Stacked bar plot showing the percentage of PC and vSMC across tissues in non-tumour (blue) and tumour (red) samples.

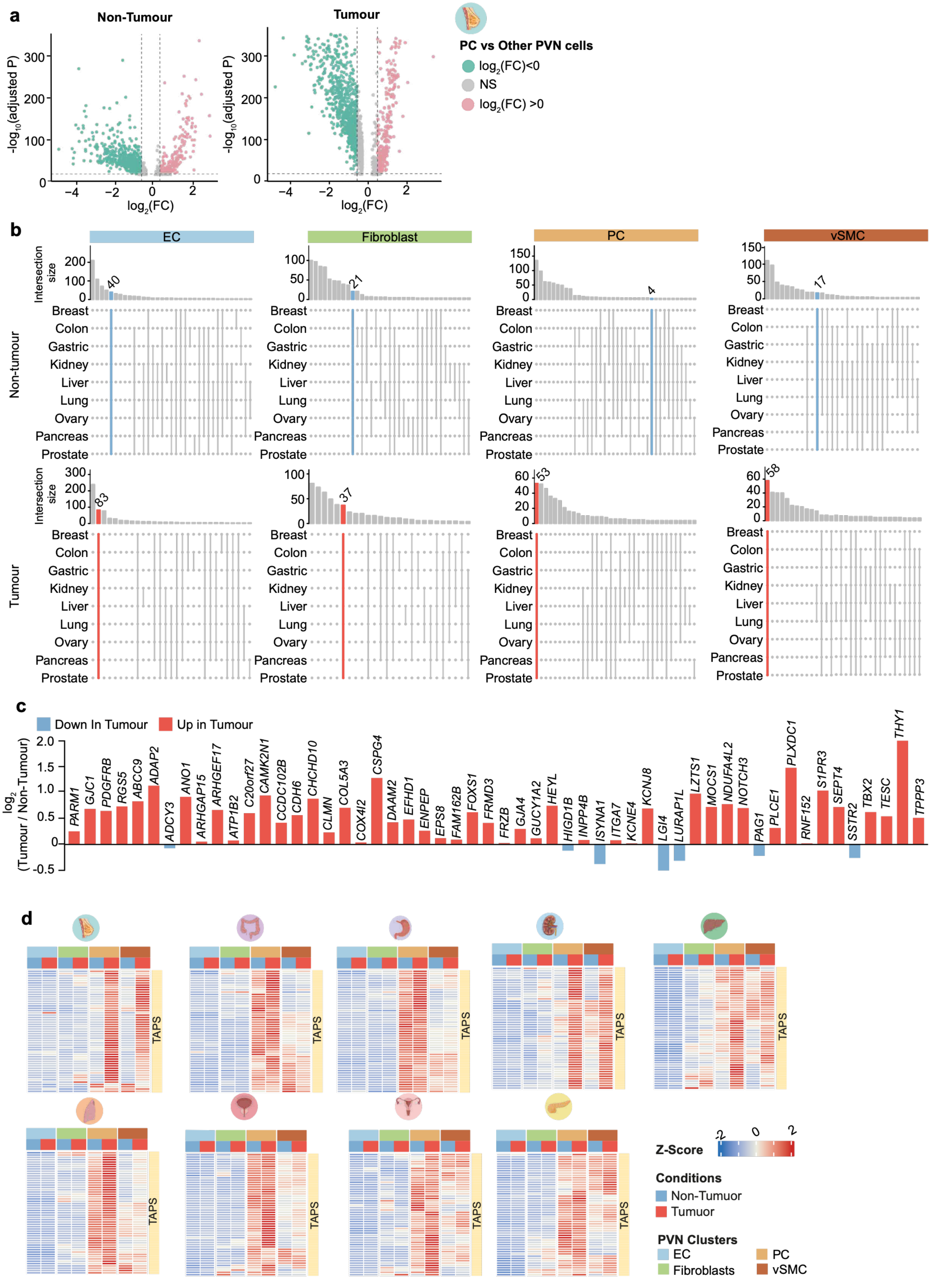

a
Non-Tumour
Tumour
-log10(adjusted P)
log2(FC)
PC vs Other PVN cells
log2(FC)<0
NS
log2(FC) >0
b
EC
Fibroblast
PC
vSMC
Intersection size
Non-tumour
Tumour
Breast
Colon
Gastric
Kidney
Liver
Lung
Ovary
Pancreas
Prostate
40
21
4
17
83
37
53
58
c
Down In Tumour
Up in Tumour
log2 (Tumour / Non-Tumour)
PARM1
GJC1
PDGFRB
RGS5
ABCC9
ADAP2
ADCY3
ANO1
ARHGAP15
ARHGEF17
ATP1B2
C20orf27
CAMK2N1
CCDC102B
CDH6
CHCHD10
CLMN
COL5A3
COX4I2
CSPG4
DAAM2
EFHD1
ENPEP
EPS8
FAM162B
FOXS1
FRMD3
FRZB
GJA4
GUCY1A2
HEYL
HIGD1B
INPP4B
ISYNA1
ITGA7
KCNE4
KCNJ8
LGI4
LURAP1L
LZTS1
MOCS1
NDUFA4L2
NOTCH3
PAG1
PLCE1
PLXDC1
RNF152
S1PR3
SEPT4
SSTR2
TBX2
TESC
THY1
TPPP3
d
TAPS
Z-Score
Conditions
Non-Tumuor
Tumuor
PVN Clusters
EC
Fibroblasts
PC
vSMC

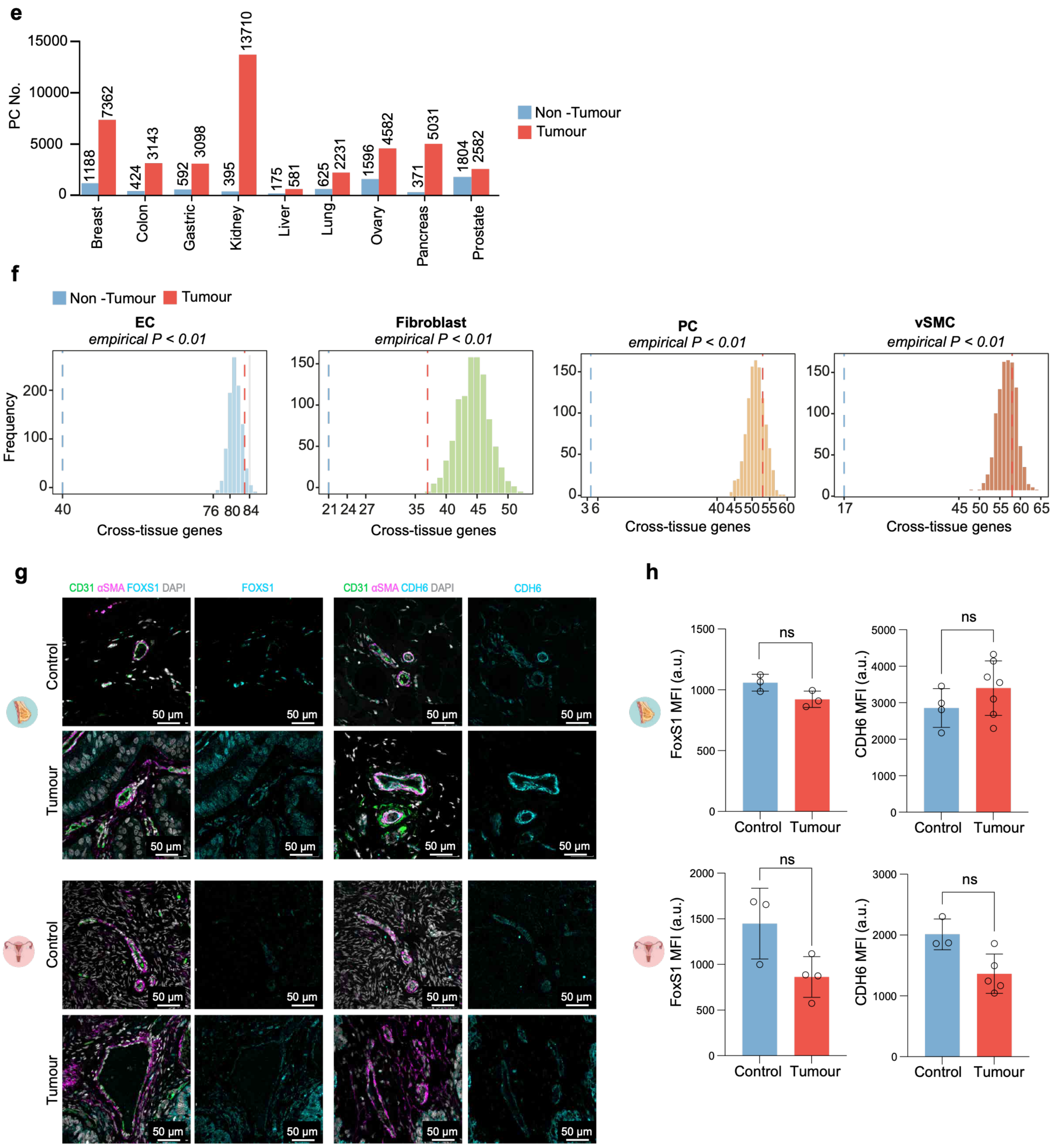

e
PC No.
Non -Tumour
Tumour
Breast
Colon
Gastric
Kidney
Liver
Lung
Ovary
Pancreas
Prostate
1188
7362
424
3143
592
3098
395
13710
175
581
625
2231
1596
4582
371
5031
1804
2582
f
EC
empirical P < 0.01
Fibroblast
empirical P < 0.01
PC
empirical P < 0.01
vSMC
empirical P < 0.01
Frequency
Cross-tissue genes
g
CD31 αSMA FOXS1 DAPI
FOXS1
CD31 αSMA CDH6 DAPI
CDH6
Control
Tumour
50 μm
h
FoxS1 MFI (a.u.)
CDH6 MFI (a.u.)
ns
Control
Tumour

**Extended Data Fig. 2:** A convergent tumour-associated transcriptional program defines pericyte identity.

**a)** Volcano plots of differentially expressed genes (DEGs) in pericytes (PC) versus other PVN populations, in the breast dataset. Comparison between non-tumour (left) and tumour (right) samples. Pink and green dots represent significantly upregulated and downregulated genes, respectively; grey dots (NS) indicate non-significantly expressed genes. **b)** UpSet plot showing the intersection of the top 250 genes across conditions in the PVN comparment. The coloured line represents genes common to all nine tissues. **c)** Bar plot showing $\log_2$(Tumour/Non-Tumour) expression of TAPS genes in pericytes. **d)** Heatmap showing the expression (Z-score) of the TAPS signature across the PVN cell types in all analysed tissues, stratified by non-tumour (blue) and tumour (red) samples. **e)** Bar plot showing the number of cells per condition within the PC compartment. **f)** Histogram representing the number of common genes identified across 1,000 iterations for each studied subtype. The blue line represents the common genes identified in each subtype (after downsampling), and the red line represents the common tumour-associated genes identified without downsampling. **g)** Representative immunofluorescence images of tumour and control human breast (top) and ovarian (bottom) tissues showing CD31+ blood vessels surrounded by αSMA+ mural cells, together with single TAPS-markers FOXS1 and CDH6. **h)** Quantification of FOXS1 and CDH6 mean fluorescence intensity (MFI) in breast and ovarian tissues. Each dot represents one patient. Data are shown as mean ± s.e.m. Statistical significance was assessed using the Mann-Whitney test; $P > 0.05$ was considered non-significant (ns).

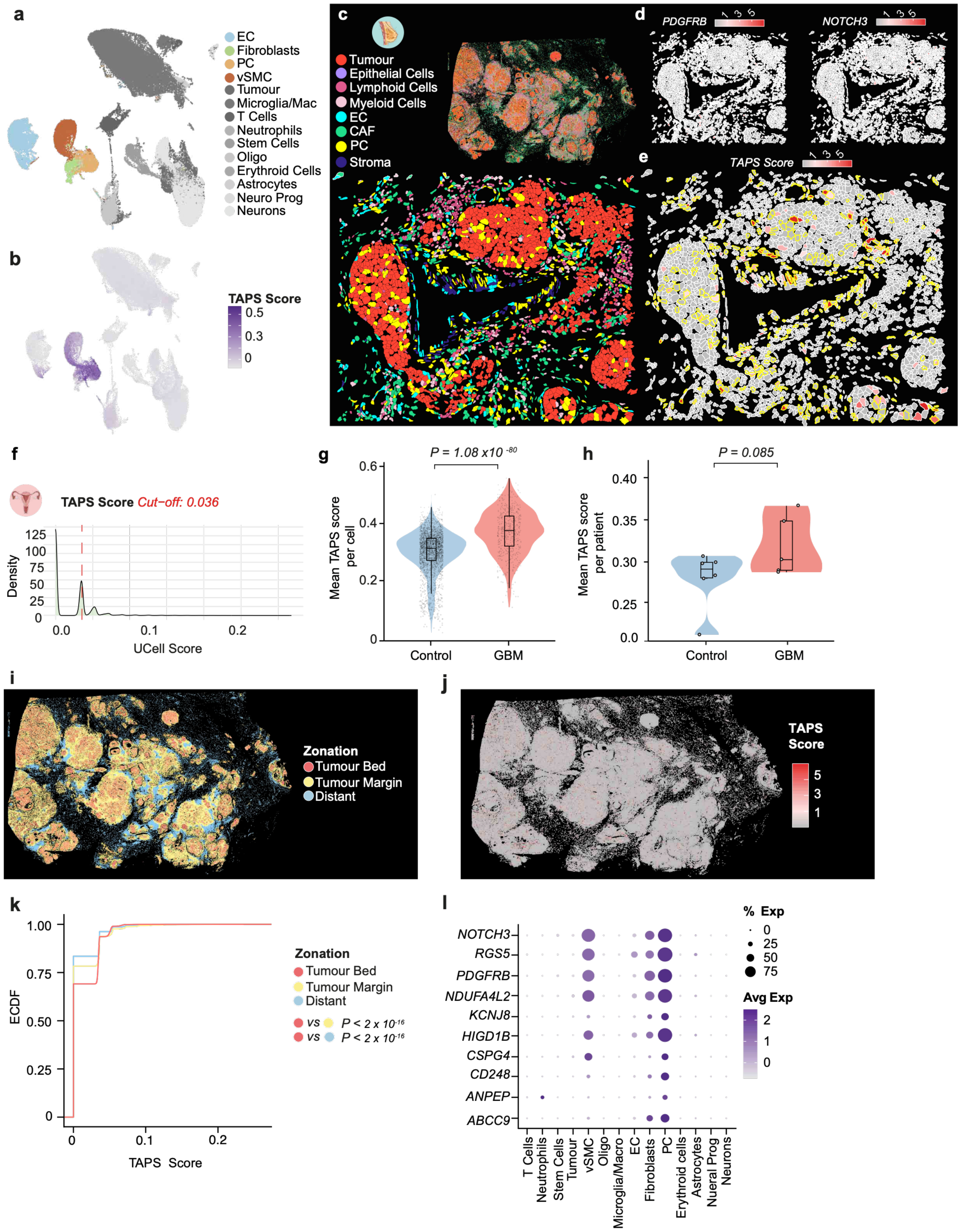

a
EC
Fibroblasts
PC
vSMC
Tumour
Microglia/Mac
T Cells
Neutrophils
Stem Cells
Oligo
Erythroid Cells
Astrocytes
Neuro Prog
Neurons
b
TAPS Score
0.5
0.3
0
c
Tumour
Epithelial Cells
Lymphoid Cells
Myeloid Cells
EC
CAF
PC
Stroma
d
PDGFRB
NOTCH3
e
TAPS Score
f
TAPS Score Cut−off: 0.036
Density
UCell Score
g
P = 1.08 x10 -80
Mean TAPS score per cell
Control
GBM
h
P = 0.085
Mean TAPS score per patient
Control
GBM
i
Zonation
Tumour Bed
Tumour Margin
Distant
j
TAPS Score
k
ECDF
TAPS Score
Zonation
Tumour Bed
Tumour Margin
Distant
vs P < 2 x 10-16
vs P < 2 x 10-16
l
NOTCH3
RGS5
PDGFRB
NDUFA4L2
KCNJ8
HIGD1B
CSPG4
CD248
ANPEP
ABCC9
% Exp
Avg Exp
T Cells
Neutrophils
Stem Cells
Tumour
vSMC
Oligo
Microglia/Macro
EC
Fibroblasts
PC
Erythroid cells
Astrocytes
Nueral Prog
Neurons

**Extended data Fig. 3.** TAPS enables robust pericyte identification across datasets and spatial contexts.

**a)** UMAP plot of the annotations provided by *Wälchli et al.* Abbreviations: Oligo, oligodendrocytes; Neuro prog, neuronal progenitors; Mac, macrophages **b)** Feature plot showing the expression of the TAPS score in the *Wälchli et al.* dataset. **c)** Spatial distribution of annotated cell types in breast. Representative images show the full tissue section (top) and a magnified inset (bottom). **d)** Representative images of the spatial expression of *PDGFRB* and *NOTCH3*. **e)** Representative image of the spatial distribution of the TAPS score; the yellow border represents pericytes (PC) identified in panel **c**. **f)** Density plot showing the distribution of TAPS scores within the ovary dataset. The red dashed line represents the 75th percentile threshold used to define TAPS-positive cells. **g)** Violin plots showing the TAPS score per cell within the PC cluster, comparing healthy brain and glioblastoma (GBM) samples. Statistics were calculated using a Wilcoxon rank-sum test. **h)** Violin plots showing the TAPS score per patient within the PC cluster, comparing healthy brain and GBM samples. Statistics were calculated using a Wilcoxon rank-sum test. **i)** Spatial distribution of the different tissue zones: tumour bed (red), tumour margin (yellow), and distant tissue (blue), in the breast sample. **j)** Spatial distribution of the TAPS UCells score across the breast sample. **k)** Empirical cumulative distribution function (ECDF) curves showing TAPS UCell scores across tumour zonation regions in breast. Statistical significance was estimated using the Kolmogorov-Smirnov test. **l)** Dot plot showing the average expression (colour scale) and percentage of expression (dot size) of canonical PC markers used to evaluate model performance in the *Wälchi et al.* dataset.

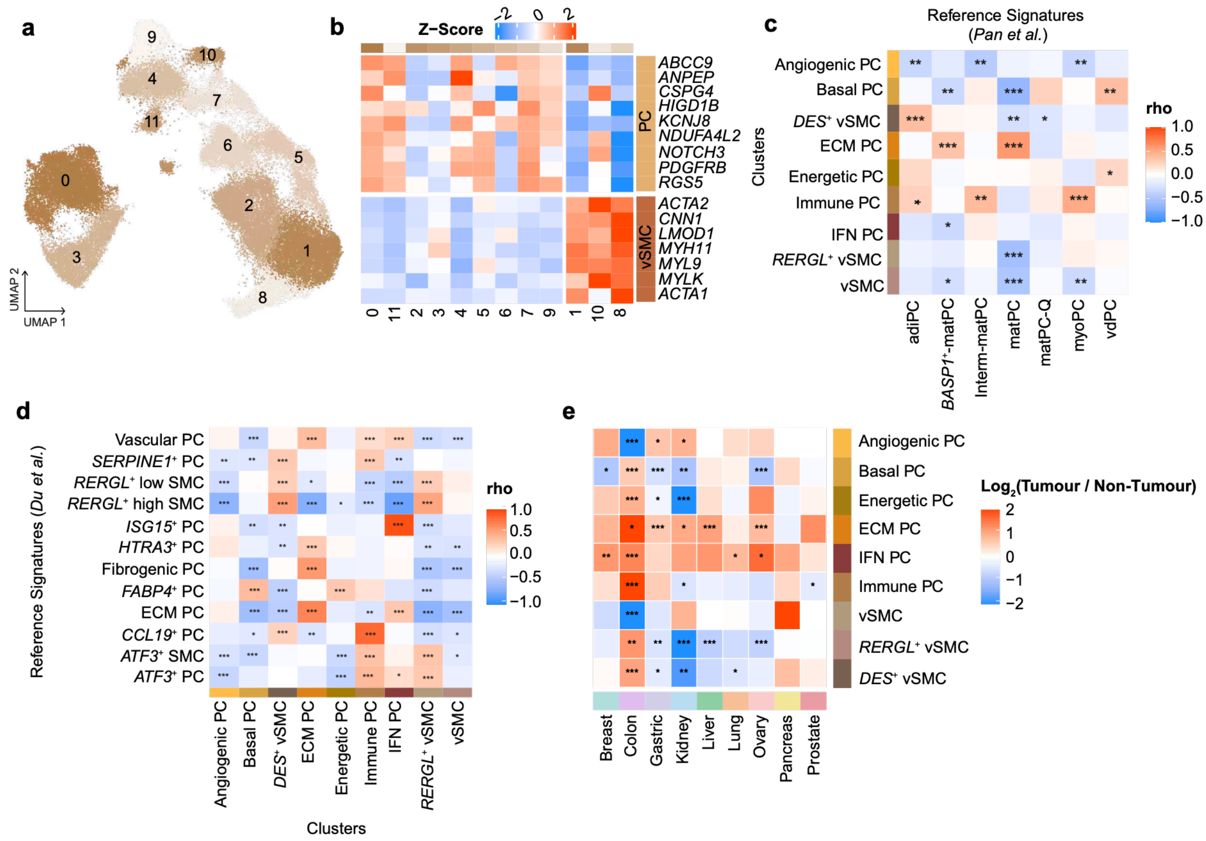


**Extended data Fig. 4.** Tumour-associated pericytes exhibit distinct functional states.
**a)** UMAP plot showing the clustering of mural cells into transcriptionally distinct subtypes. Each colour represents a specific cluster. **b)** Heatmap showing the expression patterns of canonical pericyte (PC) and vSMC marker genes. Rows correspond to marker genes and columns to mural cell subtypes. Expression values are scaled as Z-scores. **c)** Heatmap showing Spearman correlation values between the subtypes defined in this atlas and those identified in *Pan et al*. Statistical significance is indicated as follows: P ≤ 0.05 (*), P ≤ 0.01 (**), and P ≤ 0.001 (***). **d)** Heatmap showing Spearman correlation values between the subtypes defined in this atlas and those identified in *Du et al.* Statistical significance is indicated as follows: P ≤ 0.05 (*), P ≤ 0.01 (**), and P ≤ 0.001 (***). **e)** Heatmap showing the $\log_2$FC in cell proportions between tumour and non-tumour samples estimated by propeller. Positive and negative values indicate expanded and contracted populations in tumours, respectively. Tissue colour scheme as in a. Statistical significance was assessed using an empirical Bayes moderated t-test, with adjusted P values indicated as: P ≤ 0.05 (*), P ≤ 0.01 (), and P ≤ 0.001 (*).

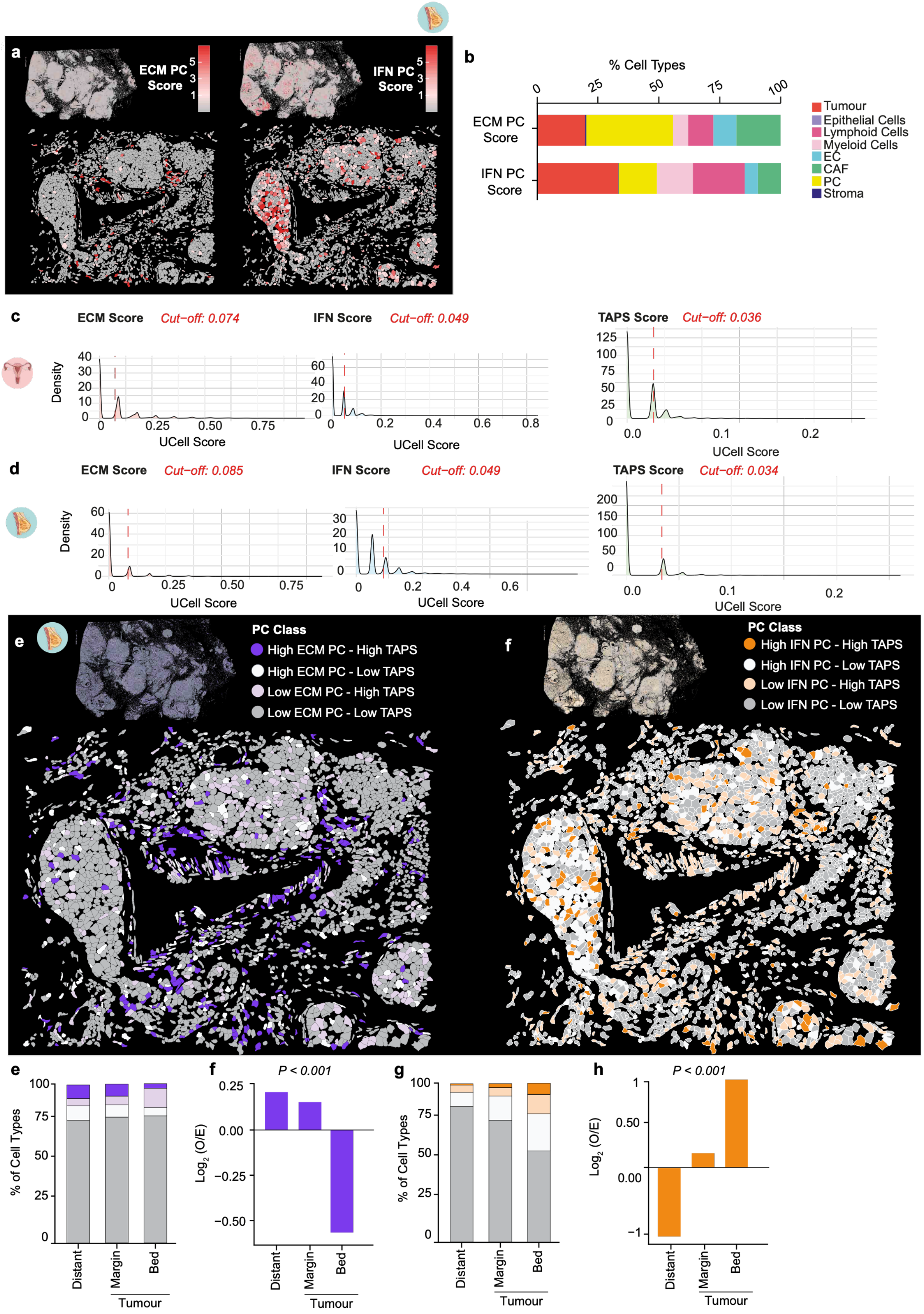

a
ECM PC Score
IFN PC Score
b
% Cell Types
0 25 50 75 100
ECM PC Score
IFN PC Score
Tumour
Epithelial Cells
Lymphoid Cells
Myeloid Cells
EC
CAF
PC
Stroma
c
ECM Score Cut−off: 0.074
IFN Score Cut−off: 0.049
TAPS Score Cut−off: 0.036
Density
UCell Score
d
ECM Score Cut−off: 0.085
IFN Score Cut−off: 0.049
TAPS Score Cut−off: 0.034
Density
UCell Score
e
PC Class
High ECM PC - High TAPS
High ECM PC - Low TAPS
Low ECM PC - High TAPS
Low ECM PC - Low TAPS
f
PC Class
High IFN PC - High TAPS
High IFN PC - Low TAPS
Low IFN PC - High TAPS
Low IFN PC - Low TAPS
e
% of Cell Types
Distant
Margin
Bed
Tumour
f
P < 0.001
Log2 (O/E)
Distant
Margin
Bed
Tumour
g
% of Cell Types
Distant
Margin
Bed
Tumour
h
P < 0.001
Log2 (O/E)
Distant
Margin
Bed
Tumour

**Extended data Fig. 5.** TAPS enables spatial resolution of specialised tumour-associated pericyte states.

**a)** Spatial distribution of ECM (left) and IFN (right) pericyte (PC) signatures, quantified by UCell scores. Representative images showing the full tissue section (top) and a magnified inset (bottom) are provided for each signature. **b)** Stacked bar plot showing the classification of cell types exhibiting positive expression for ECM and IFN signatures. **c)** Density plot of the ECM and IFN signatures scores and TAPS score across all the cell types in ovary, the red line represents 75th percentile cut off, used to determine a positive cell type. **d)** Density plot of the ECM, IFN and TAPS signatures score across all the cell types in breast, the red line represents 75th percentile cut off, used to determine a positive cell type. **e)** Spatial distribution of the ECM-associated PCs classified by the combined activity of the ECM signatures and TAPS score. Representative images showing the full tissue section (top) and a magnified inset (bottom) are provided for each signature, in breast. **f)** Spatial distribution of the IFN-responsive PCs classified by the combined activity of the IFN signature and TAPS score. Representative images showing the full tissue section (top) and a magnified inset (bottom) are provided for each signature, in breast. **e)** Stacked bar plot showing the distribution of ECM-associated PCs across the different tumour zones in breast. **f)** Bar plot showing the $log_2$-enrichment score of ECM-associated PCs across tumour zonation's in breast. Significance was determined by a chi-squared test ($P<0.01$). **g)** Stacked bar plot showing the distribution of IFN-responsive PCs across the different tumour zones in breast. **h)** Bar plot showing the $log_2$-enrichment score of IFN-responsive PCs across tumour zonation's in breast. Significance was determined by a chi-squared test ($P<0.01$).

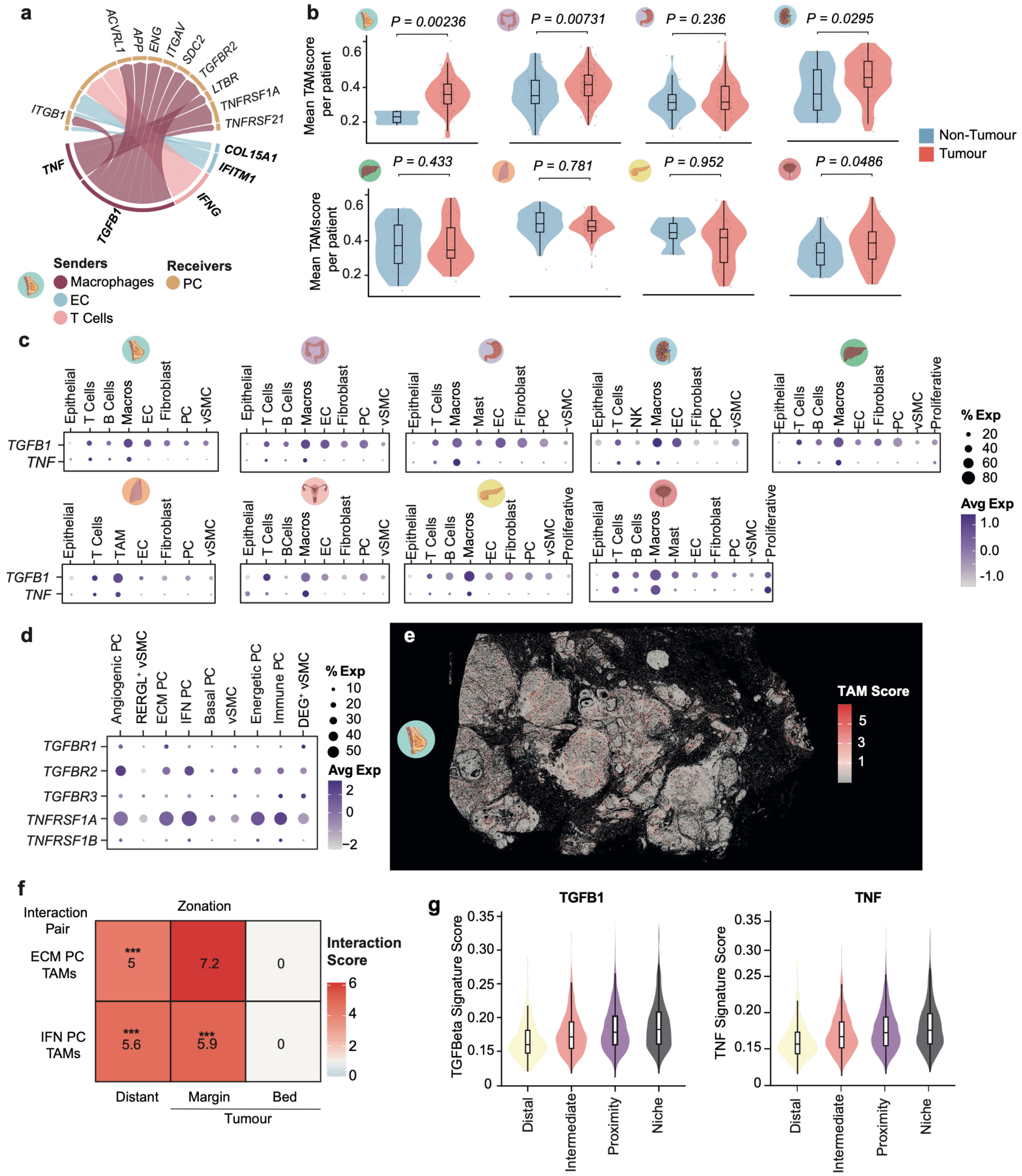
a
ACVRL1
APP
ENG
ITGAV
SDC2
TGFBR2
LTBR
TNFRSF1A
TNFRSF21
COL15A1
IFITM1
IFNG
TGFB1
TNF
ITGB1
Senders
Macrophages
EC
T Cells
Receivers
PC
b
P = 0.00236
P = 0.00731
P = 0.236
P = 0.0295
P = 0.433
P = 0.781
P = 0.952
P = 0.0486
Mean TAMscore per patient
Non-Tumour
Tumour
c
Epithelial
T Cells
B Cells
Macros
EC
Fibroblast
PC
vSMC
Mast
NK
Proliferative
TAM
BCells
TGFB1
TNF
% Exp
Avg Exp
d
Angiogenic PC
RERGL+ vSMC
ECM PC
IFN PC
Basal PC
vSMC
Energetic PC
Immune PC
DEG+ vSMC
TGFBR1
TGFBR2
TGFBR3
TNFRSF1A
TNFRSF1B
% Exp
Avg Exp
e
TAM Score
f
Interaction Pair
Zonation
ECM PC TAMs
IFN PC TAMs
Distant
Margin
Bed
Tumour
Interaction Score
g
TGFB1
TNF
TGFBeta Signature Score
TNF Signature Score
Distal
Intermediate
Proximity
Niche

**Extended data Fig. 6.** Tumour-associated macrophages shape specialised pericyte states.
**a)** Circle plot illustrating the top ligands and their corresponding receptors in breast. **b)** Violin plot showing the mean UCell score per patient for the TAM signature within macrophage populations, comparing non-tumour and tumour samples. Statistical significance was determined using a Wilcoxon rank-sum test. Tissues organized alphabetically: breast, colon, gastric, kidney, liver, lung, pancreas, and prostate. **c)** Dot plot showing the average expression (colour intensity) and percentage of expression (dot size) of the ligands *TGFB1* and *TNF* across TME populations per tissue, organized alphabetically: breast, colon, gastric, kidney, liver, lung, ovary, pancreas, and prostate. **d)** Dot plot showing the average expression (colour intensity) and percentage of expression (dot size) of *TGFBR1*,*TGFBR2*,*TGFBR3* and *TNFRSF1A*, *TNFRSF1B* receptors within mural cell subtypes. **e)** Spatial distribution of TAM signatures quantified by UCell scores; representative images show the breast tissue overview. **f)** Heatmap of interaction scores between ECM-associated and IFN-responsive PCs with TAMs across tissue zones, in breast. Statistical significance was assessed using a permutation test; FDR-adjusted P values are indicated as: adjusted P ≤ 0.05 (*), adjusted P ≤ 0.01 (), and adjusted P ≤ 0.001 (*). **g)** Violin plot with the average expression of *TGFB* (left) and *TNF* (right) target gene signatures, categorized by distance between PC and TAMs: niche (<20 μm), proximity (20-50 μm), intermediate (50-100 μm), and distal (>100 μm).

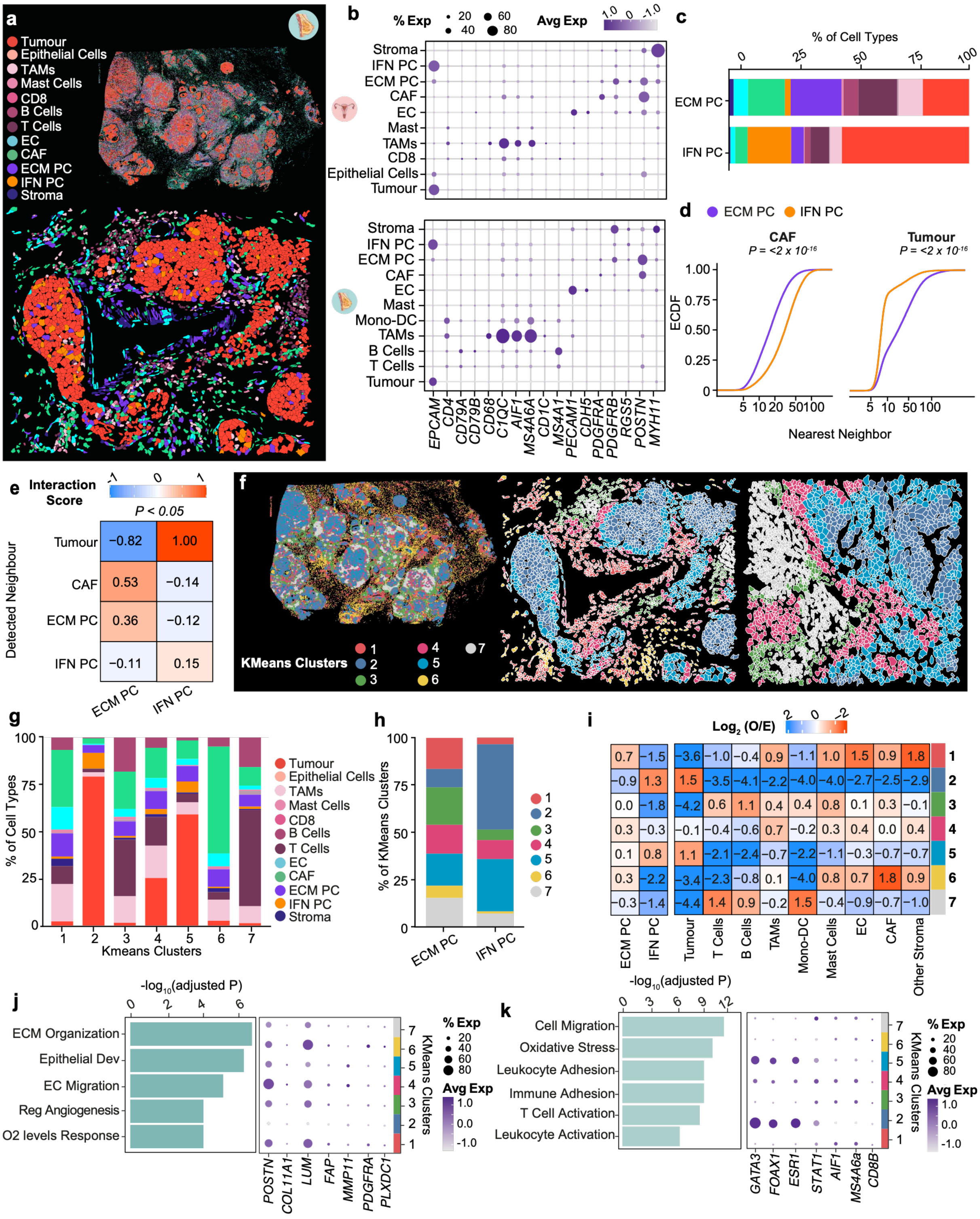

a
Tumour
Epithelial Cells
TAMs
Mast Cells
CD8
B Cells
T Cells
EC
CAF
ECM PC
IFN PC
Stroma
b
% Exp
20
40
60
80
Avg Exp
1.0
0
-1.0
Stroma
IFN PC
ECM PC
CAF
EC
Mast
TAMs
CD8
Epithelial Cells
Tumour
Mono-DC
B Cells
T Cells
EPCAM
CD4
CD79A
CD79B
CD68
C1QC
AIF1
MS4A6A
CD1C
MS4A1
PECAM1
CDH5
PDGFRA
PDGFRB
RGS5
POSTN
MYH11
c
% of Cell Types
0
25
50
75
100
ECM PC
IFN PC
d
ECM PC
IFN PC
CAF
Tumour
P = <2 x 10-16
ECDF
1.00
0.75
0.50
0.25
0
5 10 20 50 100
Nearest Neighbor
e
Interaction Score
P < 0.05
Detected Neighbour
Tumour −0.82 1.00
CAF 0.53 −0.14
ECM PC 0.36 −0.12
IFN PC −0.11 0.15
ECM PC
IFN PC
f
KMeans Clusters
1 2 3 4 5 6 7
g
% of Cell Types
Kmeans Clusters
h
% of KMeans Clusters
ECM PC
IFN PC
i
Log2 (O/E)
1: 0.7 −1.5 −3.6 −1.0 −0.4 0.9 −1.1 1.0 1.5 0.9 1.8
2: −0.9 1.3 1.5 −3.5 −4.1 −2.2 −4.0 −4.0 −2.7 −2.5 −2.9
3: 0.0 −1.8 −4.2 0.6 1.1 0.4 0.4 0.8 0.1 0.3 −0.1
4: 0.3 −0.3 −0.1 −0.4 −0.6 0.7 −0.2 0.3 0.4 0.0 0.4
5: 0.1 0.8 1.1 −2.1 −2.4 −0.7 −2.2 −1.1 −0.3 −0.7 −0.7
6: 0.3 −2.2 −3.4 −2.3 −0.8 0.1 −4.0 0.8 0.7 1.8 0.9
7: −0.3 −1.4 −4.4 1.4 0.9 −0.2 1.5 −0.4 −0.9 −0.7 −1.0
ECM PC
IFN PC
Tumour
T Cells
B Cells
TAMs
Mono-DC
Mast Cells
EC
CAF
Other Stroma
j
-log10(adjusted P)
ECM Organization
Epithelial Dev
EC Migration
Reg Angiogenesis
O2 levels Response
POSTN
COL11A1
LUM
FAP
MMP11
PDGFRA
PLXDC1
KMeans Clusters
k
Cell Migration
Oxidative Stress
Leukocyte Adhesion
Immune Adhesion
T Cell Activation
Leukocyte Activation
GATA3
FOXA1
ESR1
STAT1
AIF1
MS4A6a
CD8B

**Extended data Fig. 7.** Distinct tumour ecosystems are associated with specialised pericyte states.

**a)** Spatial distribution of the refined TME populations. Representative images showing the full tissue section (top) and a magnified inset (bottom) are provided in breast. **b)** Dot plot showing the average expression (colour intensity) and percentage of expression (dot size) of markers used to define TME populations in ovary (top) and breast (bottom). **c)** Stacked bar plot showing the distribution of TME populations surrounding ECM-associated and IFN-responsive pericytes (PC) in breast. **d)** Empirical cumulative distribution function (ECDF) curves showing the distances between ECM-associated and IFN-responsive PCs in breast. Statistical significance was estimated using the Kolmogorov-Smirnov test. **e)** Heatmap of interaction scores between ECM-associated and IFN-responsive PCs with respect to tumour cells, CAFs, and other PCs in breast. Statistical significance was estimated using a permutation-based test. **f)** K-means spatial clustering representation. Representative images showing the full tissue section (left) and two magnified insets (middle and right) displaying spatial clusters in breast. **g)** Stacked bar plot showing the distribution of TME cell types within the different defined K-means clusters in breast. **h)** Stacked bar plot showing the distribution of ECM-associated and IFN-representative pericytes across the K-means clusters in breast. **i)** Heatmap of the enrichment of the different cell types within the defined K-means clusters in breast. **j)** Bar plot of overrepresented GO terms in clusters enriched in ECM-associated PCs (clusters 7 and 3, left) and dot plot showing the average expression (colour intensity) and percentage of expression (dot size) of the top marker genes across the different clusters (right) in breast samples. **k)** Bar plot of overrepresented GO terms in clusters associated in IFN-responsive PCs (cluster 4, left) and dot plot showing the average expression (colour intensity) and percentage of expression (dot size) of the top marker genes across the different clusters (right) in breast samples.
